# Ancient and recent riverine gene flow contributed to the adaptive radiation of sailfin silversides in Wallace’s Dreampond

**DOI:** 10.1101/2025.04.16.648611

**Authors:** E L R De Keyzer, F Herder, A Böhne, F Campuzano Jiménez, V Burskaia, S Kukowka, A Tracey, A Denton, G Oatley, Wellcome Sanger Institute Tree of Life programme, Wellcome Sanger Institute Scientific Operations: DNA Pipelines collective, Tree of Life Core Informatics collective, D F Mokodongan, D Wowor, H Svardal

## Abstract

While adaptive radiations significantly contribute to the world’s biodiversity, much is unknown about the genetic and ecological factors underlying these rapid successions of speciation. It has been suggested that hybridisation can facilitate the speciation process by generating genetic diversity on which diversifying selection can act. Sailfin silverside fishes (Telmatherinidae) in the Malili Lakes system in Sulawesi have diversified within the last 2 million years. We assembled and annotated a chromosome-scale reference genome of the riverine sailfin silverside *Telmatherina bonti* and generated whole genome sequences of all species of *Telmatherina* in Lake Matano, South Sulawesi, Indonesia, one of the world’s oldest and deepest lakes. We reconstructed the phylogenetic relationships within the adaptive radiation of sailfin silversides and inferred past and ongoing introgression patterns. Genome-wide tests confirmed two monophyletic clades, sharpfins and roundfins. However, within clades, we found mismatches between morphology-based taxonomic assignments and genome-wide genetic relationships. We found signs of both old and ongoing introgression between river-dwelling *T. bonti* and the lacustrine sharpfin group, as shown in elevated D-statistic, f4-ratio and f-branch statistic. Levels of excess allele sharing between riverine species and the three most common lacustrine species declined with increasing distance from the river-inlet, indicating ongoing introgression at the lake-river interface. This combination of past and ongoing hybridisation in a radiating species flock makes Lake Matano *Telmatherina* a particularly valuable system to study fundamental mechanisms driving rapid speciation under genomic exchange. The phylogenomic framework elaborated in this study provides the foundation for studies of the processes shaping this charismatic radiation.

## Introduction

An important aspect of biodiversity is how new species arise and establish. Adaptive radiations, rapid bursts of ecological speciation originating from a small number of ancestral lineages, have significantly contributed to the world’s biodiversity (Berner and Salzburger 2015). The molecular mechanisms underlying these rapid speciation events are still largely unclear. Specifically, it is not well known what genetic and ecological factors underlie the difference between radiating lineages and lineages which do not diversify despite apparent ecological opportunity. Understanding phylogenomic relationships and patterns of evolutionary history of an adaptive radiation opens opportunities to study mechanisms of speciation within the radiation.

The genomic diversity required for rapid speciation can be obtained through introgressive hybridization with a related lineage (Seehausen 2004; Arnold 2006; Nolte and Tautz 2010). This was for example the case in Darwin’s finches, for which introgressive hybridization is the major factor contributing to genomic diversity (Lamichhaney et al. 2015). Hybridisation has often been detected within young adaptive radiations (Malinsky et al. 2018; Kozak et al. 2021). Although genetic exchange can break down reproductive isolation, potentially slowing down or reversing speciation (Seehausen et al. 2008), recent studies emphasise its potential role as a source of genetic variation. Introgression can even give rise to novel traits, increasing adaptive potential and thereby facilitating speciation (Seehausen 2004; Seehausen 2013; Marques et al. 2019; Peñalba et al. 2024). Due to the complexity of the evolutionary mechanisms underlying adaptive radiation, there are still many unknown factors about the molecular basis of adaptive radiation and natural systems in which the effects of ongoing genetic exchange can be studied are rare. To get a clear understanding of the genomic makeup and especially the role of introgression in an ongoing radiation, ideally these processes need to be studied in an ecologically well documented system with relatively smaller complexity compared to the larger and often interconnected adaptive radiations such as East African cichlid fishes or Heliconius butterflies.

The Malili Lakes in Central Sulawesi, Indonesia, were termed Wallace’s Dreamponds (Herder, Nolte, et al. 2006) for their status as a hotspot of aquatic biodiversity, with adaptive radiations and high degrees of endemism in snails, shrimp, crabs, and fishes (Vaillant et al. 2011; von Rintelen et al. 2012). Sailfin silverside fishes (Telmatherinidae) have diversified in the lakes within the last 1-2 million years (Stelbrink et al. 2014). Previous studies suggest 19 distinct species (of which 15 are formally described) with distinctive phenotypic and ecological adaptations (Herder, Schwarzer, et al. 2006). In Lake Matano, the most upstream and deepest lake of the Malili Lakes system, an adaptive radiation of sailfin silversides has produced species that conquered a diverse set of ecological niches (Herder et al. 2008; Pfaender et al. 2016). Lake Matano sailfin silversides can be attributed to two ecologically and morphologically well-defined groups distinguished by the shape of the second dorsal and the anal fin (Kottelat 1991), (Herder, Nolte, et al. 2006); the bentho-pelagic roundfins and the predominantly benthic sharpfins (Herder, Schwarzer, et al. 2006), shown to be genetically distinct based on AFLP markers (Herder and Schliewen 2010). Diversification within the roundfin and sharpfin lineages had been dated to approximately 1.0 and 0.9 million years ago (Mya), respectively, using mitochondrial markers (Stelbrink et al. 2014). Both clades have diversified and inhabit a variety of ecological niches, with corresponding morphological adaptations related to resource use, including variations in body shape and size, oral and pharyngeal jaw structure and the length of gill rakers (Pfaender et al. 2010; Wasiljew et al. 2021). Morphological differentiation is complex, covering different structures rather than single key innovations (Pfaender et al. 2010).

Two species of roundfins have formally been described: *T. prognatha* Kottelat, 1991, and *T. antoniae* Kottelat, 1991 (Kottelat 1991), yet large differences in morphology, AFLP markers, trophic ecology, habitat use as well as the observation of assortative mating suggest *T. antoniae* consists of two separate species (Herder et al. 2008; Pfaender et al. 2011; Wasiljew et al. 2021), referred to as *T. antoniae* ‘small’ and *T. antoniae* ‘large’. Roundfin *Telmatherina* have, to different degrees, a pelagic ecology, contrasted to the predominantly epibenthic sharpfins (Herder et al 2008). The sharpfin species flock is the most diverse lineage of sailfin silversides in terms of adaptive traits (Herder, Schwarzer, et al. 2006; Pfaender et al. 2010; Pfaender et al. 2016). The six reported species of sharpfins (of which four are formally described) (Herder, Schwarzer, et al. 2006), have mainly been distinguished by morphometric and meristic typing (Kottelat 1991). Trophic groups partly coincide with the species groups, and include insect (*T. opudi* Kottelat, 1991), shrimp (*T.* sp. ‘thicklip’) and mollusc (*T. wahjui* Kottelat, 1991) feeders, fish predators (*T.* sp. ‘elongated’*, T. abendanoni* Weber, 1913) and a specialised fish egg predator (*T. sarasinorum* Kottelat, 1991) (Pfaender et al. 2010; Pfaender et al. 2016). Sharpfins show species-specific behaviour, often related to feeding strategies (Cerwenka et al. 2012) and mating (Gray and McKinnon 2006). To some extent, a correlation between genotypic and phenotypic variation has been observed (Schwarzer et al. 2008; Pfaender et al. 2016), but previous genetic investigations have been limited to AFLP and mitochondrial data. At the same time, trophic niches of sharpfin species widely overlap and there is substantial overlap across species in morphological variation (Pfaender et al. 2016), indicative of a young, evolving species flock (Herder et al. 2008; Pfaender et al. 2010).

Although the majority of morphological and species diversity within *Telmatherina* is represented by lacustrine species, riverine populations are common in the rivers around the Malili lakes (Herder, Schwarzer, et al. 2006). *T. wahjui* from the outlet river Petea and the area in Lake Matano around Petea river displays physical and behavioural adaptations to a riverine habitat, such as a wider head and adaptations in mating behaviour (Gray and McKinnon 2006). Riverine populations of *T.* cf. *bonti* occur in the streams surrounding and between the Malili Lakes (Herder, Schwarzer, et al. 2006; Herder, Nolte, et al. 2006). Finally, *T. bonti* Weber & de Beaufort, 1922 has been reported from Lawa river, the inlet stream of Lake Matano (Hadiaty and Wirjoatmodjo 2002). *T. bonti* are ram feeders, specialised on obtaining small food items in their fast-flowing stream habitats (Wasiljew et al. 2022).

Reproductive isolation among different species of sailfin silversides is substantial (Herder et al 2006a, 2008), yet there are signals of ongoing gene flow between some species (Herder, Nolte, et al. 2006; Herder et al. 2008; Schwarzer et al. 2008; Stelbrink et al. 2014). Previous marker-based efforts failed to completely resolve phylogenetic relationships between *Telmatherina* species and mixed signals were interpreted as gene flow both within sharpfins and within roundfins (Herder et al. 2008; Stelbrink et al. 2014). There are indications of high levels of introgression between lacustrine and riverine lineages (Herder, Nolte, et al. 2006; Schwarzer et al. 2008), including between the riverine *T. wahjui* population in river Petea and lacustrine sharpfins in Lake Matano (Schwarzer et al. 2008). Sharpfins were found to be monophyletic based on AFLP data (Herder, Nolte, et al. 2006), yet they cluster into two mitochondrial haplogroups (Stelbrink et al. 2014). While some individuals carry haplotypes with a sister group relationship to the lake’s roundfins, others carry mitochondria related to riverine lineages. This signal was interpreted as riverine introgression early in the divergence of sharpfin silversides (Herder, Nolte, et al. 2006; Stelbrink et al. 2014), or even ongoing hybridization (Schwarzer et al. 2008). However, given the limited number of investigated markers this data is not sufficient to provide a conclusive picture. A better understanding of the complete evolutionary history of these species will require expanding information through obtaining high resolution genome wide markers.

Disentangling the evolutionary history of the adaptive radiation of sailfin silversides, including past and ongoing introgression, is an important step towards developing this system as a model to study speciation mechanisms. In this study we assembled a reference genome of the riverine *T. bonti* and investigated the phylogenomic relationships and patterns of past and ongoing introgression between sailfin silversides (*Telmatherina*) from Lake Matano, Sulawesi and from stream populations. Using whole genome sequences of 36 specimens attributed to eleven species of sailfin silversides, we tested for signals of past introgression from riverine species into Lake Matano sharpfins, relative to their sister group roundfins, and presumed hybridisation on ecological timescales between riverine and lake species. Our analysis reveals a complex evolutionary history of Lake Matano *Telmatherina* and implicates a role of both ancient and ongoing introgression in shaping and maintaining current genetic diversity.

## Results

### High quality reference genome and annotation for *Telmatherina bonti*

A high-quality reference genome of *T. bonti* (fTelBon1.1, GCA_933228915.1) was constructed as part of the tree of life project (https://www.sanger.ac.uk/programme/tree-of-life/) (supplementary figure S1, supplementary figure S2). The final assembly had a total length of 986,039,709 bp in 146 sequence scaffolds with scaffold N50 and N90 sequence lengths of 41,523,244 and 35,388,860 bp respectively (fig. 2a). 99.6% of the assembly sequence was assigned to 25 chromosomal-level scaffolds, representing 24 autosomes, numbered by sequence length, and the mitochondrion (Supplementary table S1). The assembly had a BUSCO completeness of 98% using the actinopteryggi_odb10 reference set (fig. 2b). The Ensembl gene annotation system (Aken et al. 2016) was used to generate an annotation for the fTelBon1.1 assembly. The resulting annotation included 47,104 transcripts assigned to 21,413 coding and 4,664 non-coding genes.

### Genome data supports roundfin and sharpfin clades and most taxa

To investigate whether previous morphology-based taxonomic assignments are consistent with genome-wide genetic relationships, we analysed whole genome sequences of 36 *Telmatherina* specimens from Lake Matano (fig. 1, supplementary figure S3, Supplementary table S2). We constructed a neighbour joining (NJ) tree based on genome-wide pairwise differences (supplementary figure S4). In addition, we built genome-wide and mitochondrial maximum-likelihood (ML) phylogenies and performed PCA (fig. 3-4). All methods showed a clear separation between the bentho-pelagic roundfins and the predominantly epibenthic sharpfins. Within roundfins we found clear support for the previously identified ecomorphological groups *T. prognatha*, *T. antoniae* ‘small’, and *T. antoniae* ‘large’, which each formed monophyletic groups in the ML consensus tree and the genome-wide NJ tree (fig. 3a, supplementary figure S4) and well-defined clusters in a PC analysis only including roundfin samples (supplementary figure S5). We found similarly strong support for three sharpfin species *T.* sp. ‘elongated’, *T.* sp. ‘thicklip’, and *T. wahjui* (fig. 3c, fig. 4a-c). Conversely, samples of the most common sharpfin species in Lake Matano, *T. sarasinorum*, *T. opudi*, and *T. abendanoni*, did not show any consistent pattern of grouping by taxonomic assignment in any of the phylogenetic trees (fig. 2, supplementary figure S4) or PC analyses (fig. 4b-c). Instead, those individuals showed a relatively large spread intermediate between *T.* sp. ‘thicklip’ and *T.* sp. ‘elongated’ along PC2 of an analysis only including sharpfin samples (fig. 4c).

**Fig. 1.**
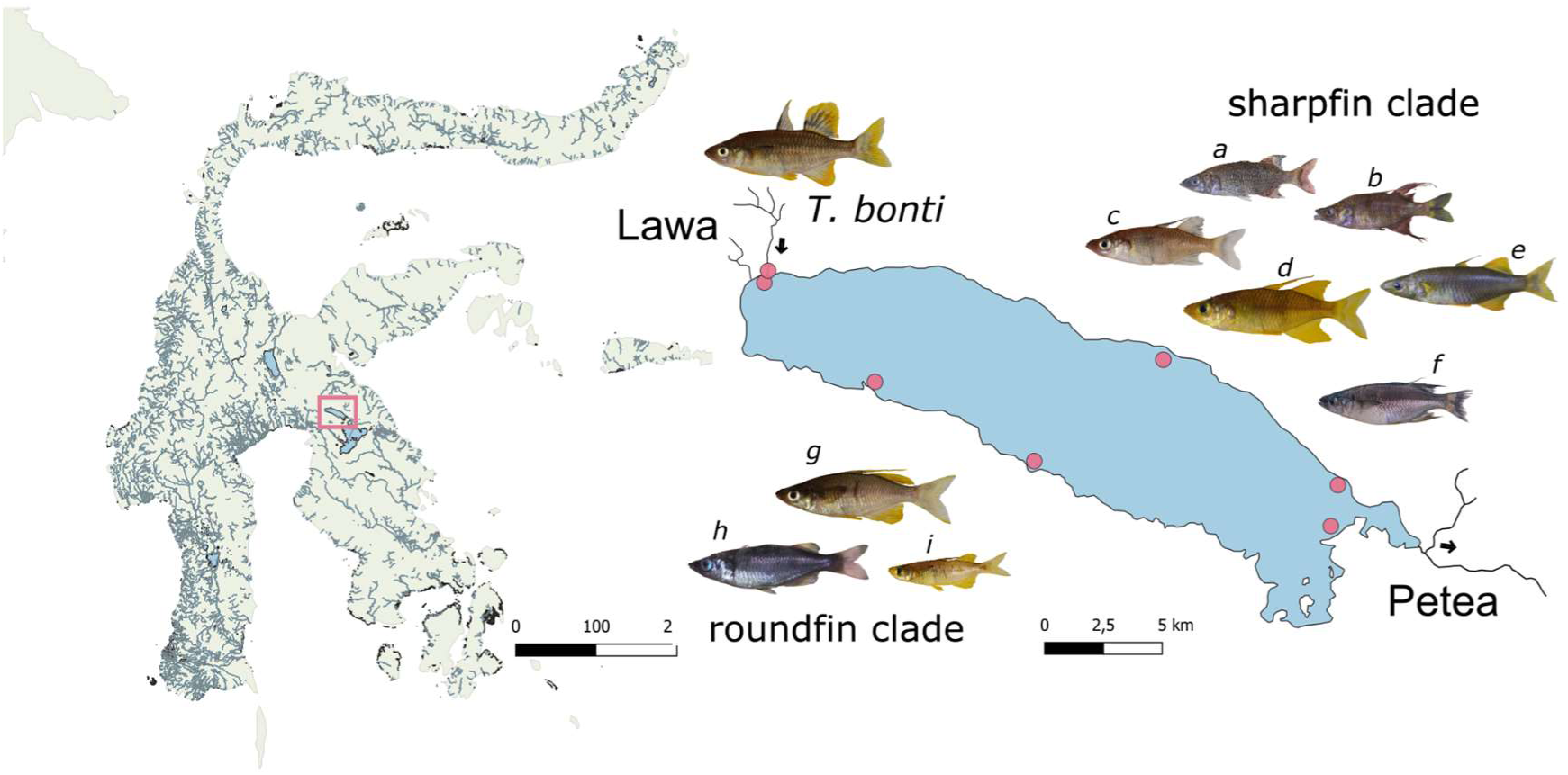
Map of sampling locations. Location of Lake Matano in the Malili Lakes system, Sulawesi (left). Detail of lake Matano and the inflow river Lawa and outflow river Petea (right). Dots represent the six sampling sites along Lake Matano and one in the inflow Lawa river. Pictures depict the six species of sharpfin Telmatherina (*a T. abendanoni* (n = 3)*, b T.* sp. ‘thicklip’ (n = 3)*, c T. opudi* (n = 6)*, d T. sarasinorum* (n = 6)*, e T.* sp. ‘elongated’ (n = 3)*, f T. wahjui* (n = 3)) and three species of roundfin Telmatherina *(g T. antoniae* ‘large’ (n = 3), *h T. prognatha* (n = 3)*, i T. antoniae* ‘small’ (n = 3)) and the riverine *T. bonti* (n = 3).

**Fig. 2.**
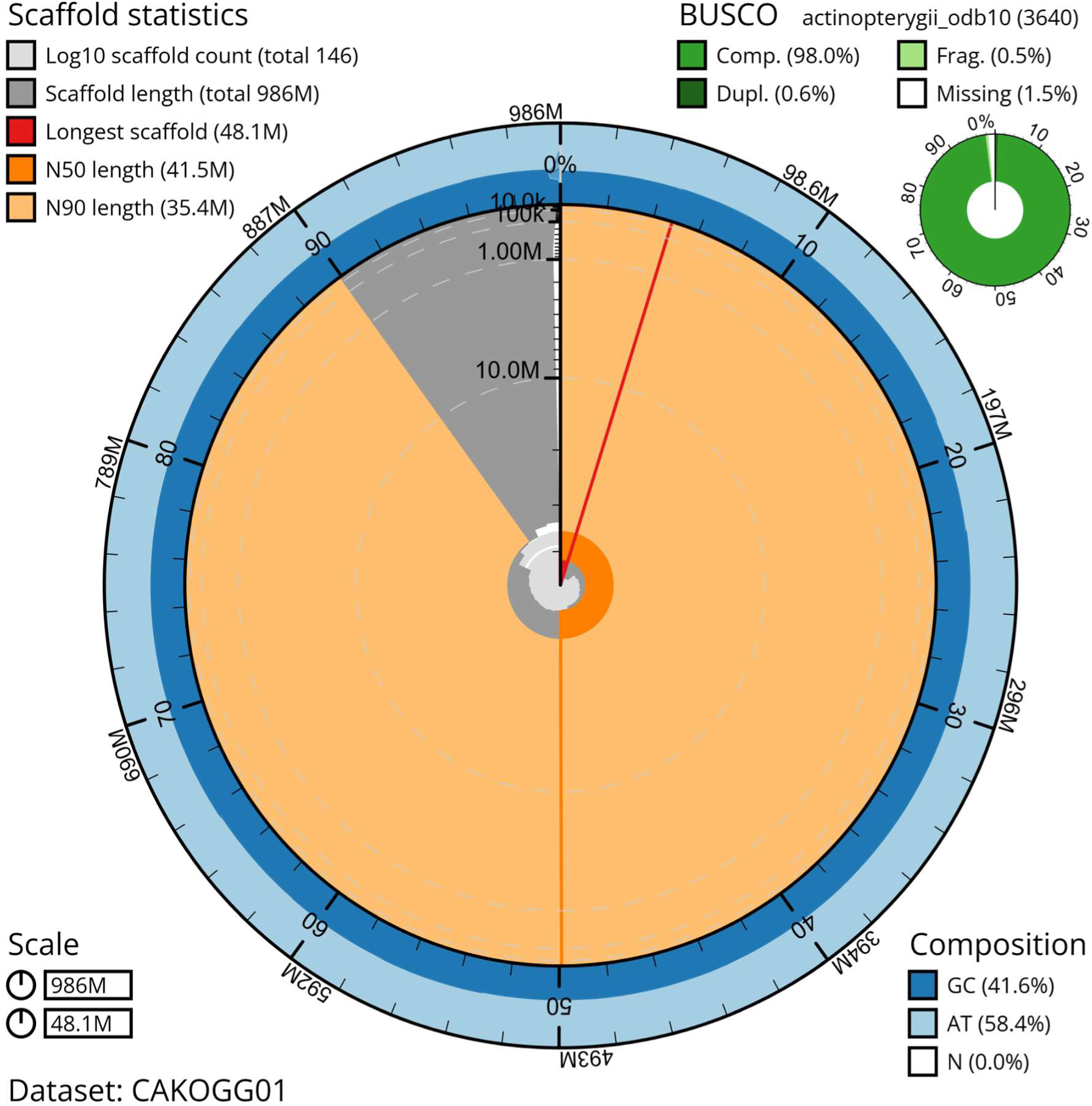
Assembly statistics for reference genome assembly of *Telmatherina bonti*. The BlobToolKit Snailplot shows N50 metrics and BUSCO gene completeness. The main plot is divided into 1,000 size-ordered bins around the circumference with each bin representing 0.1% of the 986,039,709 bp assembly. The distribution of sequence lengths is shown in dark grey (length measured from outside to inside) with the plot radius scaled to the longest sequence present in the assembly shown in red (48,058,054 bp). Orange and pale-orange arcs show the N50 and N90 sequence lengths (41,523,244 and 35,388,860 bp), respectively. The pale grey spiral shows the cumulative sequence count on a log scale with white scale lines showing successive orders of magnitude. The blue and pale-blue area around the outside of the plot shows the distribution of GC, AT and N percentages in the same bins as the inner plot. Top right: summary of complete, fragmented, duplicated and missing BUSCO genes in the actinopterygii_odb10 set. An interactive version of this figure is available at https://blobtoolkit.genomehubs.org/view/Telmatherina%20bonti/dataset/CAKOGG01/snail.

**Fig. 3.**
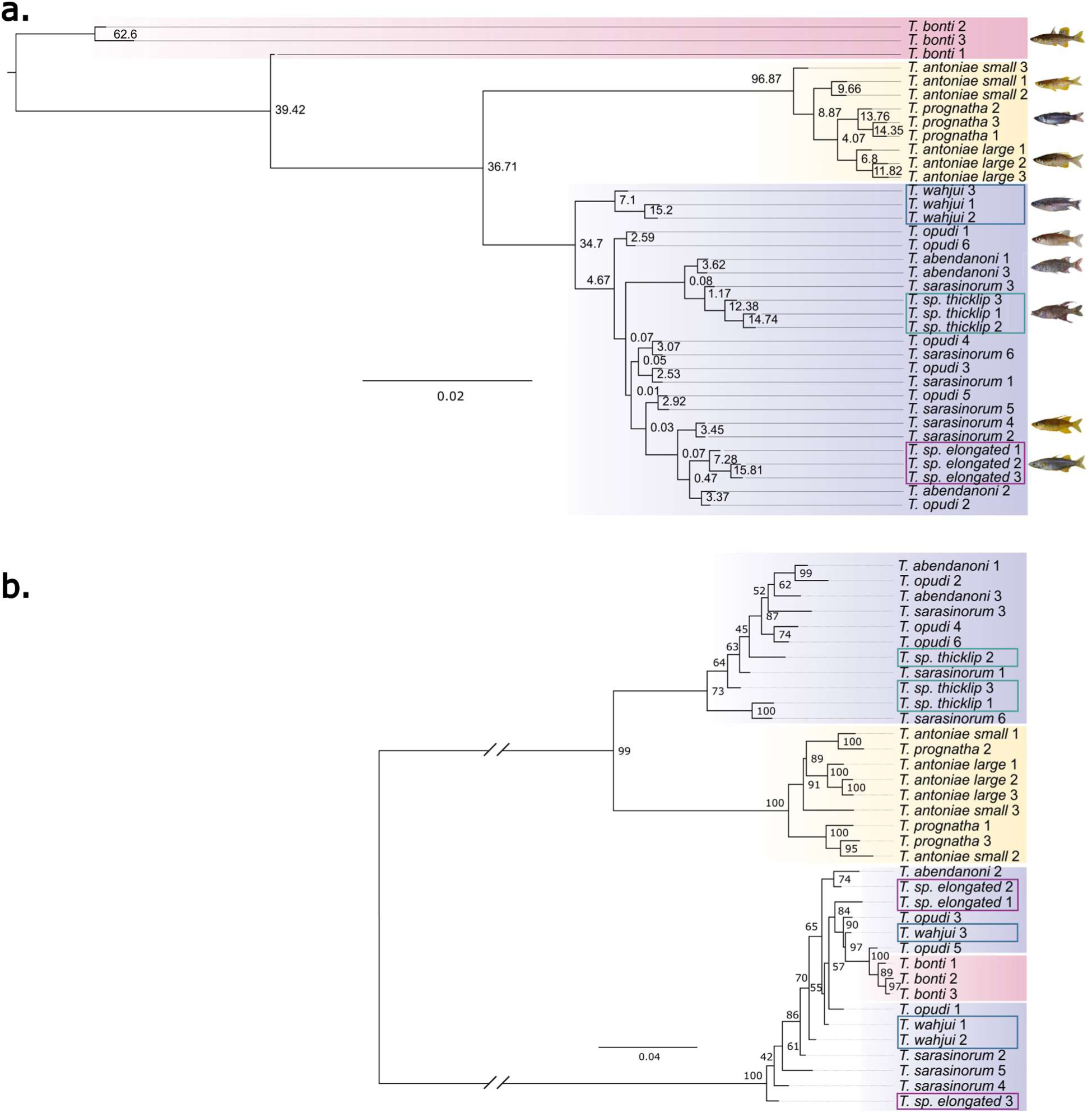
Phylogeny based on maximum likelihood (ML) trees. Sharpfin species are highlighted in purple, roundfins in yellow and the *T. bonti* samples are indicated in red background shading. **a.** Consensus phylogeny based on 9150 ML trees in 100 kb windows across the genome. Numbers indicate the proportion of windows supporting this split. **b.** Phylogeny based on mitochondrial DNA. Numbers indicate bootstrap support for 1000 bootstrap replicates. Coloured boxes highlight the positions of the three sharpfin species monophyletic in the genome-wide tree shown in panel a.

**Fig. 4.**
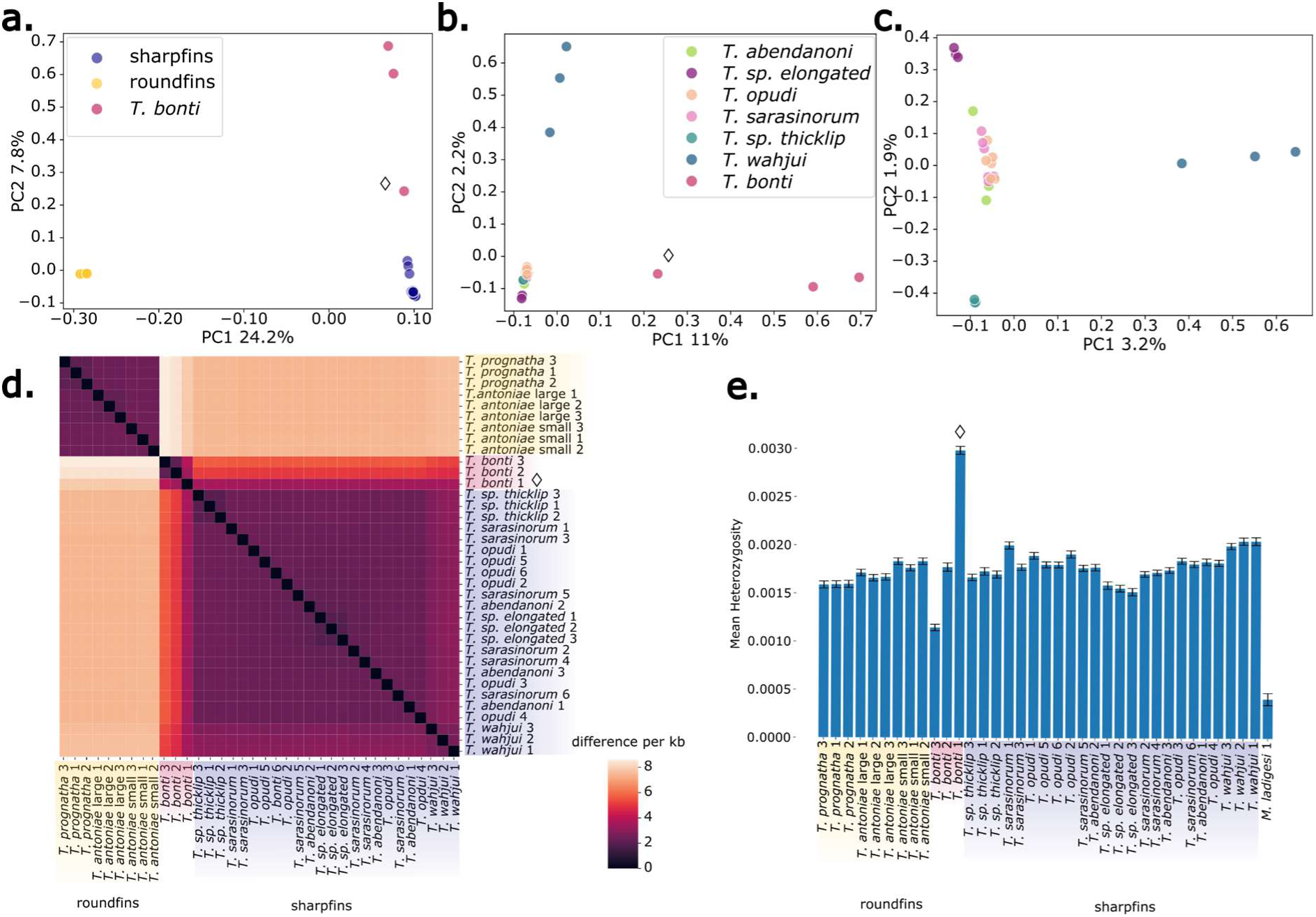
Principal component analysis and pairwise sequence divergence. **a-c**. Visualisation of the first and second genomic principal components. **a.** shows the Lake Matano sharpfins and roundfins, and the riverine *T. bonti*. **b**. Lake Matano sharpfins separated by species and *T. bonti* **c.** only Lake Matano sharpfin species. **d.** Pairwise sequence divergence between individual samples visualised in a heatmap. **e.** Mean heterozygosity in individual samples calculated over 200kb windows, error bars indicate 95% confidence interval between windows. Diamond shape (◊) indicates *T. bonti* 1, the presumed recent hybrid sample in panels a, b, d and e.

### Complex relationships between lake and river dwelling species

Our sample set contained members of two stream-dwelling species: *T. wahjui* from Petea river, the outflow of Lake Matano, and *T. bonti* from Lawa river, the major tributary of Lake Matano. As mentioned above, *T. wahjui* clearly belongs to the sharpfin genetic clade, in which they fall basally to all other sharpfins (fig. 3a). That said, there was a relatively large spread of these individuals in the PC analysis of sharpfins (fig. 4b) consistent with variable ancestry proportions. *T. bonti*, the second stream dwelling species, showed an even more extreme spread among samples in the PC analysis, albeit along different axes of variation (PC2 in the analysis of all samples, fig. 4a; PC1 in an analysis of only sharpfins and *T. bonti*, fig. 4b). Consistent with this, *T. bonti* samples also showed inconsistent phylogenetic clustering for different inference methods (fig. 3, supplementary figure S4). Specifically, while in the genome-wide neighbour joining tree (supplementary figure S4) the three samples formed a sister group to the sharpfins, they formed an outgroup to both sharpfins and roundfins in the ML consensus tree (fig. 3a). An astral consensus of NJ based local distance trees, on the other hand, gave an intermediate pattern with two *T. bonti* samples clustering as a global outgroup and one as a sister group to sharpfins (supplementary figure S4). Together, these analyses suggest that river-dwelling *Telmatherina* show within-population variation in their genetic relationships with the lacustrine species groups.

### Mitochondrial phylogeny splits into three haplotype groups

Our sample set was divided into three groups based on mitochondrial haplotypes (fig. 3b). One group, consisting entirely of roundfin samples, was in a sister relationship with a group that included approximately half of the sharpfin samples. The third group contained the remaining sharpfin samples along with all *T. bonti* samples. A similar pattern was reported by Stelbrink et al. (2016), who referred to the roundfin and sharpfin haplotypes in the sister group as “the original Matano haplotype”. All samples of the stream-dwelling sharpfin species *T. wahjui* and the lacustrine predator *T. sp.* ‘elongated’ shared a haplotype similar to *T. bonti*. In contrast, all samples of the shrimp-eating sharpfin species *T. sp.* ‘thicklip’ carried the original sharpfin haplotype. The remaining three sharpfin species—*T. opudi*, *T. sarasinorum*, and *T. abendanoni*—did not consistently group with either haplotype cluster: roughly half of the samples had a *T. bonti*-like haplotype, while the others carried the original sharpfin haplotype. None of the roundfin nor sharpfin species were monophyletic in their haplotype groups.

### Incomplete lineage sorting and low variation within clades

Given the relatively young age of the radiation, the incomplete sorting of standing genetic variation between speciation events (incomplete lineage sorting, ILS), could lead to variation in genetic relationships among species along the genome. And indeed, we found that among 9150 maximum likelihood trees, based on non-overlapping genomic windows of 100kb, there was substantial topological variation. Consistent with this we found allele sharing across all groups (Supplementary figure S6). Pairwise distance was lowest among roundfin samples and highest between roundfin and *T. bonti* and sharpfin samples (fig. 4d).

### Signals of introgression between lake and river dwelling species

To identify genetic introgression among lake species and between lake and river dwelling species, we tested for signals of excess allele sharing. Elevated D-statistic, f4-ratio and f-branch statistics point towards introgression between *T. bonti* and all the sharpfin species (fig. 5a, supplementary figure S7, supplementary figure S8) and to a lesser extent the roundfin species, and between *T. wahjui* and other sharpfin species. Comparison of admixture proportions of *T. bonti* into the sharpfins *T. abendanoni*, *T. sarasinorum* and *T. opudi* from different sampling sites revealed a spatial gradient of declining admixture with increasing distance from the river-lake interface (fig. 5c-d). This is suggestive of relatively recent gene flow having happened at the river-lake interface that is diffusing into lake sharpfin populations through migration. One of the samples in our dataset, “*T. bonti* 1”, while morphologically identified as *T. bonti*, appears to be a recent *T. bonti* x sharpfin hybrid based on its position in the phylogenetic trees (fig. 3a), in the PCA (fig. 4c), and its high level of heterozygosity (fig. 4e).

**Fig. 5.**
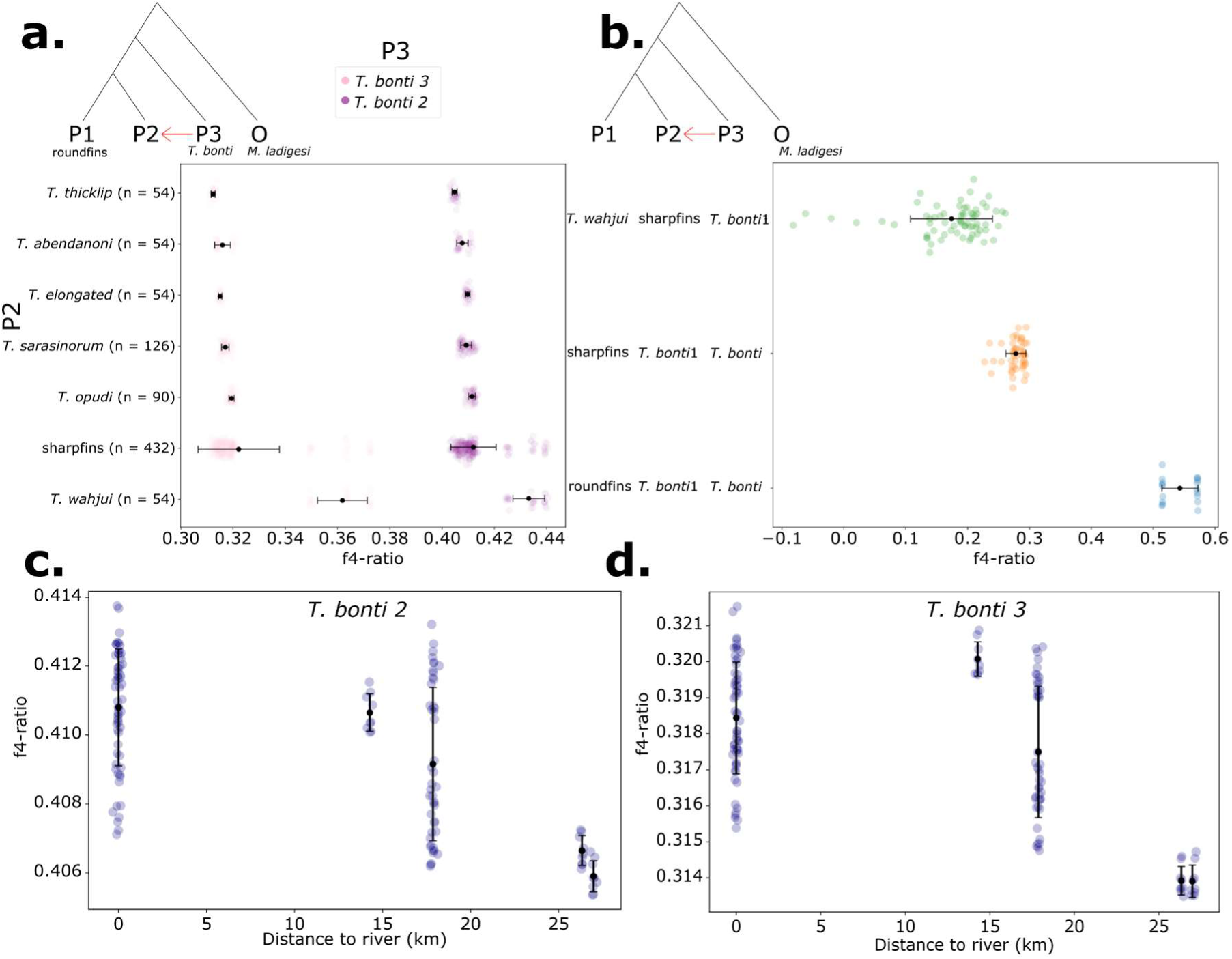
Introgression statistics. **a.** Excess allele sharing (f4-ratio) between *T. bonti* 3 (pink; P3) and *T. bonti* 2 (purple; P3) and the sharpfin species (P2), relative to the roundfins (P1). **b**. Excess allele sharing (f4-ratio) between different groups, P1, P2 and P3 are indicated on the Y-axis. *T. bonti* 1 is the presumed recent hybrid individual, *T. bonti* are all other *T. bonti* samples. Higher values indicate more introgression from P3 into P2, relative to P1 **c-d.** excess allele sharing (f4-ratio) of *T. bonti* 2 (**c**) and *T. bonti* 3 (**d**) (P3) into three sharpfin species (*T. sarasinorum*, *T. opudi*, *T. abendanoni*; P2) compared to roundfins (P1) at different distances from the river-lake interface. Higher values indicate more introgression from *T. bonti* into the three sharpfin species.

## Discussion

Using whole genome sequences, we reconstructed the phylogenomic relationships and ancient and recent introgression history within the adaptive radiation of *Telmatherina*. We find that morphology-based taxonomic assignments are not consistent with genome-wide genetic relationships for three of the lacustrine species, *T. opudi*, *T. sarasinorum* and *T. abendanoni*. In addition, river-dwelling *Telmatherina* show within-population variation in their genetic relationships with the lacustrine species groups. We confirm a mito-nuclear discordance in the sharpfin clade, indicative of introgression between sharpfins and *T. bonti*. Introgression between riverine species and the lacustrine *Telmatherina* is apparent in elevated levels of excess allele sharing, which decline with increasing distance from the lake-river interface.

### New species status for sharpfins and roundfins

Our results confirm a phylogenomic separation between roundfins and sharpfins, both are monophyletic clades within the Lake Matano sailfin silversides. Our results support the sharpfins *T. wahjui*, *T.* sp. ‘thicklip’, and *T.* sp. ‘elongated’ as separate taxonomic entities, as suggested by previous work based on morphology, behaviour and diet (Herder, Schwarzer, et al. 2006), supporting formal taxonomic descriptions of *T.* sp. ‘thicklip’ and *T.* sp. ‘elongated’. A fourth group of sharpfins consists of all samples of *T. opudi*, *T. sarasinorum* and *T. abendanoni*, which together form one cluster in PCA, but are inconsistently placed in phylogenetic trees. While previous research shows morphological and ecological variation between these species (Herder, Schwarzer, et al. 2006; Pfaender et al. 2016), we could not differentiate between them in our genome wide analysis. Possible interpretations include ongoing hybridisation between these three species (Pfaender et al. 2016), hybridisation with a species outside of the sharpfin clade, distorting the phylogenomic signal within the clade, or an absence of separate gene pools.

The phylogenetic signal for the roundfin clade is clear throughout our analysis and consistent with the described morphospecies, *T. prognatha*, *T. antoniae* ‘small’ and *T. antoniae* ‘large’ (Herder et al. 2008; Pfaender et al. 2011). This result extends previous studies using fewer genetic markers, which found no (Herder, Nolte, et al. 2006) or only low (Herder et al. 2008) genetic differentiation between the three roundfin species. Based on whole genome sequences, we find *T. antoniae* ‘large’ to be more closely related to *T. prognatha* than to *T. antoniae* ‘small’. We consider *T. antoniae* ‘large’ and *T. antoniae* ‘small’ as separate species, and urge for an official taxonomic revision of the roundfin clade, based on our findings regarding the genomic phylogeny and previous findings regarding morphology, adaptive niche differentiation, assortative mating, and diet (Herder et al. 2008; Pfaender et al. 2011). The species status of *T. prognatha* has not been under debate in the literature and does also not raise any concerns in our analysis. Even though the overall phylogenetic signal in the roundfins is clear, we found alternative topologies where roundfin species are not monophyletic in some parts of the genome, and the maximum likelihood tree based on mitochondrial DNA does not clearly distinguish between the three roundfin species. This inconsistency could indicate that reproductive isolation between the species is recent or be a signal of incomplete isolation.

Similar discordance between morphological and ecological species definitions and genomic signals had been reported for example for *Coenonympha* butterflies (Greenwood et al. 2025) and *Eudesmia* trees (McLay et al. 2023). This discordance had been attributed to amongst others incomplete lineage sorting and interspecific gene flow, complicating genomic species delineation. The fact that our genomic investigation revealed discordance between described species and genetic groups observed in Lake Matano *Telmatherina*, calls for an investigation of species hypotheses throughout the broader silverside radiation, including the *Telmatherina* and closely related *Paratherina* and *Tominanga* of the other lakes in the Malili system and nearby lakes.

### Speciation and signatures of past introgression in *Telmatherina*

In this study, we found strong signals of past introgression of the riverine *T. bonti* into lacustrine sharpfins, shown as elevated values of the f4-ratio and f-branch statistic. We speculate that this introgression might have played a role in increasing diversity of the sharpfin clade, by providing access to a broader range of alleles for selection to act upon. Indeed, the lacustrine *Telmatherina* show an increased number of trophic traits and colour variation compared to other families of endemic Lake Matano fishes (Pfaender et al., 2010, 2016). In addition, of the two clades of *Telmatherina* in Lake Matano, the sharpfin clade, which is more species rich and shows higher variation in morphology, behaviour and trophic adaptations (Roy et al. 2004; Gray and McKinnon 2006; Pfaender et al. 2010; Cerwenka et al. 2012), is also the clade which received most introgression from *T. bonti.* Thus, our results offer a preliminary indication based on whole genome data that, as has been suggested before (Herder, Nolte, et al. 2006; Schwarzer et al. 2008), the evolutionary history of sharpfin *Telmatherina* has in part been shaped by introgression from a lineage related to the riverine *T. bonti* into an ancestor of the sharpfin clade. Similar processes have been suggested in cichlids, where gene flow between lake species and river species played an important role in exchanging genetic diversity, potentially catalysing speciation in the lake populations (Loh et al. 2013; Berner and Salzburger 2015). Ancient hybridization at the base of the Malawi cichlid radiation led to elevated divergence between splits in the radiation due to the persisting genetic variation (Svardal et al. 2020). Hybridization between two divergent lineages and multiple subsequent hybridisation events facilitated diversification in the Lake Victoria Region Superflock of Haplochromine cichlid fishes into over 700 diverse species which arose in a timespan of 150.000 years (Meier et al. 2017; Meier et al. 2023). Furthermore, it was shown recently that differentiation between two Heliconius species was driven by introgression of ecological traits (Rosser et al. 2024). Detailed assessment of divergence patterns and introgression patterns in *Telmatherina* will provide insight into the effect of gene flow on diversity in the sharpfin clade.

Overall, most species of sharpfins have similar levels of excess allele sharing with *T. bonti*, suggesting that introgression of *T. bonti* into the sharpfin clade happened long ago, either early into the sharpfin lineage or into one of the sharpfin species after which hybridisation between sharpfins homogenised introgressed alleles. In contrast, the sharpfin species *T. wahjui*, which is confined to river Petea and the lake area close to its outflow, and has morphological and behavioural adaptations to the riverine environment (Gray and McKinnon 2006; Schwarzer et al. 2008), shows more introgression from *T. bonti* compared to other sharpfin species. We speculate that the current population of *T. wahjui* is the result of past intermixing between a *T. bonti*-like lineage in the outflow river Petea and the lacustrine sharpfins. This way *T. wahjui* genetically clusters with sharpfins, while retaining genomic and phenotypic building blocks from its *T. bonti-*like ancestral genetic backbone. This interpretation is supported by our finding that the recent sharpfin-bonti hybrid (*T. bonti* 1) in our dataset shares fewer alleles with *T. wahjui* compared to the other sharpfins, in contrast to the other *T. bonti* samples. Alternatively, *T. wahjui* may have received *T. bonti* introgression in the same ancient introgression event as the rest of the sharpfin species and may have retained more of the *T. bonti* alleles, as these would be more adaptive in their riverine habitat. A thorough timing of hybridisation events and insight into the selection processes in *T. wahjui* will shed light on these questions.

### Recent and ongoing gene flow between Matano sharpfins and the riverine group

Besides signals of past gene flow, we also detected recent gene flow between the stream-dwelling *T. bonti* and Matano species. In the maximum likelihood (ML) consensus trees, assuming a GTR model, *T. bonti* is positioned as an outgroup to both the sharpfins and roundfins. On the other hand, the NJ tree suggests closer genetic similarity between *T. bonti* and sharpfins relative to roundfins, possibly due to reduced genetic distance resulting from recent gene flow. While the NJ method reflects genetic distance, not accounting for speciation models, ML is more robust for complex evolutionary histories, including variable substitution rates and heterogeneity across the genome. Given these strengths, we interpret the ML tree as more likely reflecting true phylogeny. In addition to inconsistency between NJ and ML phylogenies, we also find that mitochondrial and nuclear patterns are discordant. While the nuclear trees show monophyletic groups for sharpfins and roundfins, in the mitochondrial tree, sharpfins are split between a group that is a sister group to roundfins, and a more basal group which also contains the riverine *T. bonti*. This latter group contains besides *T. bonti,* all samples of *T. wahjui* and *T.* sp. ‘elongated’, and about half of the samples of *T. sarasinorum, T. opudi* and *T. abendanoni*. These results suggest gene flow from the riverine *T. bonti* into most species of the sharpfin clade. Using whole genome data, our study confirms the patterns identified in previous studies based on AFLP markers (Herder, Nolte, et al. 2006; Stelbrink et al. 2014).

One of the three samples in our dataset which were initially identified as *T. bonti* is a hybrid between *T. bonti* and a Lake Matano sharpfin species. This is evident from its position in the PCA intermediate between the other *T. bonti* samples and Lake Matano sharpfins. Heterozygosity in this individual is twice as high compared to the other samples, suggesting this to be a recent hybrid. The specimen in question was captured at the lake-river interface, where river-dwelling and lake-dwelling species are in direct contact. Since in our limited sample-set we already find a recent hybrid, this suggests that hybridisation might be a locally common process. The other two samples of *T. bonti* in our dataset have less sharpfin-like DNA, which suggests that these are backcrosses between *T. bonti* x sharpfin hybrids and *T. bonti*. This finding suggests the interesting possibility of ongoing introgression of sharpfin alleles into the riverine population, the opposite direction compared to ancient introgression, which introduced alleles from *T. bonti* into Lake Matano sharpfins. Sampling upstream Lawa river will reveal if there is a spatial gradient of proportion of introgressed sharpfin alleles in *T. bonti*.

We detected clear signals of decreased excess allele sharing (lower f4-ratios) with increasing distance to the river-lake interface between *T. bonti* and three sharpfin species which occur over the entire lake, *T. sarasinorum*, *T. opudi* and *T. abendanoni*. This pattern is consistent with a hybrid zone at the interface between the inlet river Lawa and the lake, possibly facilitated by natural selection favouring hybrid individuals in this ecologically intermediate zone. This spatial decline in excess allele sharing is a strong indication of ongoing or recent gene flow, as signals of older gene flow would over time have homogenised over the extent of the lake, erasing spatial signals.

### Signals of incomplete speciation

Our results show weak genomic clustering and weak lineage sorting in three sharpfin species; *T. sarasinorum*, *T. opudi* and *T. abendanoni*. Incomplete reproductive isolation between these species (Herder et al, 2006b, 2008, Schwarzer et al. 2008, Stelbrink et al. 2014), and the occurrence of sharpfin *Telmatherina* with morphological traits intermediate between species, suggest that the sharpfin system is in a dynamic stage of the species flock formation process, evolving under disruptive ecological selection. Previous studies found that variation in morphology between the sharpfin species was linked to variation in a small proportion (13%) of nuclear markers, and intermediate, less fit individuals were common, indicative of a young species flock (Pfaender et al. 2016). This signature of recent species flock formation is surprising since diversification within sharpfins has been estimated to have started relatively long ago, around 1 mya (Stelbrink et al. 2014). Why does sharpfin speciation remain incomplete? An interesting contributing factor could be that the influx of similar *T. bonti* alleles into the sharpfin species through the ongoing hybridisation with riverine *T. bonti* continues to homogenise the sharpfin genomes. This continuous influx of *T. bonti* alleles, combined with species-specific adaptation pressures, could create a balance between homogenisation of the genome and ecological selection, so that speciation remains incomplete. To understand the processes behind the incomplete separation of these sharpfin species, future studies will need to gain insight into the interplay between genetic influx of *T. bonti* into the sharpfin species, the ongoing mixing between the sharpfin species, and the strength and type of selection that introgressed alleles undergo.

### Conclusion

This study provides data supporting both ancient and ongoing hybridisation in the *Telmatherina* of Lake Matano, which may have accelerated adaptation to a wide variety of ecological niches. This combination of ancient and ongoing hybridisation in a comprehensive, still radiating flock makes the *Telmatherina* of Lake Matano a particularly valuable study system to unravel the fundamental mechanisms driving rapid speciation under genomic exchange. The reference genome and phylogenomic framework elaborated in this study provide the foundation for more in depth studies of this charismatic radiation.

## Methods

### Sampling and sequencing

We selected 36 samples of the genus *Telmatherina* from the ichthyology collections of the Leibniz Institute for the Analysis of Biodiversity Change, Museum Koenig in Bonn, Germany (Supplementary table S2). Samples were collected in Indonesia between 2002 and 2010, sampling was done by gillnet. Sex was determined visually upon capture, the fish were photographed and fish were euthanized by administering an overdose of MS222. A finclip was immediately stored in 97% ethanol; fish specimens were stored in formalin for several days before being transferred to ethanol. Two specimens of *Marosatherina ladigesi* were sampled from an aquarium population to serve as an outgroup in this study. These fish were euthanized by administering an overdose of MS222, photographed and a part of the right pectoral fin was stored separately in 97% ethanol. DNA for short read sequencing was extracted from fin clips using the Zymo tissue kit according to manufacturer’s guidelines. Extracted DNA was sent to BGI, Hong Kong and submitted for whole genome resequencing on DNBSEQ for 150bp paired end short read sequencing.

For the reference genome construction and annotation, one specimen of *T. bonti* from an aquarium stock at ZFMK was immediately dissected after euthanization with an overdose of MS222. Fin and muscle tissue samples were placed in pre-chilled tubes, immediately submerged in liquid nitrogen for snap freezing and kept at -80° C until high-molecular weight DNA extraction for long read sequencing. The workflow for high-molecular weight (HMW) DNA extraction at the Wellcome Sanger Institute (WSI) Tree of Life Core Laboratory includes a sequence of core procedures: sample preparation; sample homogenisation, DNA extraction, fragmentation, and clean-up. In sample preparation, the fTelBon1 sample was weighed and dissected on dry ice (Jay et al. 2023). Tissue from muscle was homogenised using a PowerMasher II tissue disruptor (Denton et al. 2023). HMW DNA was extracted using the Automated MagAttract v1 protocol (Sheerin et al. 2023). DNA was sheared into an average fragment size of 12–20 kb in a Megaruptor 3 system (Todorovic et al. 2023). Sheared DNA was purified by solid-phase reversible immobilisation (Strickland et al. 2023), using AMPure PB beads to sample to eliminate shorter fragments and concentrate the DNA. The concentration of the sheared and purified DNA was assessed using a Nanodrop spectrophotometer and Qubit Fluorometer using the Qubit dsDNA High Sensitivity Assay kit. Fragment size distribution was evaluated by running the sample on the FemtoPulse system.

RNA was extracted from muscle tissue of fTelBon1 in the Tree of Life Laboratory at the WSI using the RNA Extraction: Automated MagMax™ *mir*Vana protocol (do Amaral et al. 2023). The RNA concentration was assessed using a Nanodrop spectrophotometer and a Qubit Fluorometer using the Qubit RNA Broad-Range Assay kit. Analysis of the integrity of the RNA was done using the Agilent RNA 6000 Pico Kit and Eukaryotic Total RNA assay.

Pacific Biosciences HiFi circular consensus DNA sequencing libraries were constructed according to the manufacturers’ instructions. Poly(A) RNA-Seq libraries were constructed using the NEB Ultra II RNA Library Prep kit. DNA and RNA sequencing was performed by the Scientific Operations core at the WSI on Pacific Biosciences Sequel IIe (HiFi) and Illumina NovaSeq 6000 (RNA-Seq) instruments. Hi-C data were also generated from muscle tissue of fTelBon1 using the Arima-HiC v2 kit. The Hi-C sequencing was performed using paired-end sequencing with a read length of 150 bp on the Illumina NovaSeq 6000 instrument.

### Reference genome construction

The reference genome for *T. bonti* (GCA_933228915.1; https://www.ncbi.nlm.nih.gov/assembly/GCA_933228915.1) was sequenced from a single male *T. bonti* specimen bred at the LIB Bonn, Germany. A total of 36-fold coverage in PacBio long reads. The HiFi reads were first assembled using Hifiasm version 0.16.1-r375 (Cheng et al. 2021) with the --primary option. Haplotypic duplications were identified and removed using purge_dups version 1.2.3 (Guan et al. 2020). The Hi-C reads were mapped to the primary contigs using bwa-mem2 (Vasimuddin et al. 2019). The contigs were further scaffolded using the provided Hi-C data (Rao et al. 2014) in YaHS version yahs-1.1.91eebc2 (Zhou et al. 2023) using the --break option. The scaffolded assemblies were evaluated using Gfastats (Formenti et al. 2022), BUSCO (Manni et al. 2021) and MERQURY.FK (Rhie et al. 2020).

The mitochondrial genome was assembled using MitoHiFi version 2 (Uliano-Silva et al. 2023), which runs MitoFinder (Allio et al. 2020) and uses these annotations to select the final mitochondrial contig and to ensure the general quality of the sequence.

Manual curation was primarily conducted using PretextView 0.2.5 (Harry 2022), with additional insights provided by JBrowse2 (Diesh et al. 2023) and HiGlass (Kerpedjiev et al. 2018). Scaffolds were visually inspected and corrected as described by Howe et al. (Howe et al. 2021). Manual assembly curation corrected 207 missing joins or mis-joins and one haplotypic duplication, which reduced the scaffold number by 54.26 and increased the scaffold N50 by 11.67%. The process is documented at https://gitlab.com/wtsi-grit/rapid-curation.

In total 24 chromosome-scale scaffolds were confirmed by the Hi-C data, which have been named in order of size (large to small). Besides the 24 chromosomes and the mitochondrial genome, the genome assembly includes 115 unplaced scaffolds.

### Short read quality control and mapping

Whole genome sequencing reads were filtered by BGI using the SOAPnuke software (Chen et al. 2018). Sequencing reads which matched > 25% adapter sequence, had a read length of < 150 bp, > 0.1% N content or a Phred score < 33 were filtered out. Sequencing yielded ∼80.1 million clean reads (after filtering) per sample. Sequences were quality checked using FastQC v0.11.9 (Andrews 2010) and MultiQC v1.12 (Ewels et al. 2016). Reads were aligned against the *T. bonti* (NCBI:txid446457) reference genome fTelBon1.1 with BWA-MEM (BWA version 0.7.17, (Li 2013)). The resulting SAM files were compressed to cram format and quality checked using samtools flagstat. After filtering and mapping each sample to the *T. bonti* reference genome, the resulting effective sequencing coverage varied between 13.8-22.9 fold, except for the outgroup samples (*M. ladigesi*) which yielded on average 10.5 fold coverage, related to their larger distance to the reference species.

Only single nucleotide polymorphisms (SNPs) were retained for genetic investigation. Variant sites were called jointly for all samples with the mpileup function in BCFtools v1.14 (Danecek et al. 2021). Total depth and mean coverage were calculated per site. Any sites with an overall mapping quality Phred-scaled score of less than 50 (4.9% of SNPs) or with more than 10% of mapped reads with a mapping quality zero (4.0% of SNPs) were masked. Additionally, filters were applied to sites with excess heterozygosity (InbreedingCoeff < 0.2; 0.03% of SNPs), before filtering and sites with a Phred-scaled p-value of binomial test for allele balance violation of > 40 (0.06% of SNPs). Sites with > 40% missing genotypes (2.0% of SNPs) as well as sites where the sum of overall depth for all samples was unusually high (> 3 units of standard deviation above individual mean coverage; 3.6% of SNPs) were filtered out. In total, 5.2% of SNPs were removed from the dataset (Supplementary figure S9), note that most sites were removed due to multiple filters. After filtering, 43.862.995 biallelic SNPs across 24 chromosomes remained. The mitogenome of the *T. bonti* reference was 16.560 bp long. Filters were applied to mitochondrial sites with a mapping quality Phred-scaled score under 10 (19% of SNPs), sites with more than 50% of mapped reads with a mapping quality of zero (11% of SNPs), sites with a Phred-scaled p-value of binomial test for allele balance violation of > 40 (1.2% of SNPs), and sites with > 40% missing genotypes after application of other filters (23% of SNPs)(Supplementary figure S9). After filtering, 1836 SNPs were retained in the aligned mitogenome.

### Phylogeny

#### Nuclear genome

To infer evolutionary relationships between the different *Telmatherina* species we constructed genomic phylogenies based on whole genome sequences. The SNP dataset was split into genomic windows of 100kb based on SNP coordinates, using a custom python script, resulting in 9831 windows. Of these, 681 windows containing fewer than 10 SNPs were removed since these did not contain sufficient information to construct a reliable tree, resulting in a final dataset of 9150 windows, containing 10 - 7905 SNPs (median = 5793.5).

In these remaining genomic windows of 100 kb over the nuclear genome, local neighbour joining (NJ) trees were calculated based on distance matrices and a consensus tree was calculated with Astral-III (version 5.7.1, (Zhang et al. 2018)). For the same dataset, Maximum likelihood (ML) trees were inferred using the General time reversible (GTR) model in IQ-TREE v2.2.0 (Minh et al. 2020), which accounts for unequal substitution rates between nucleotide pairs and unequal base frequencies. Bootstrap support was calculated for 1000 bootstrap replicates with ultrafast bootstrap approximation (UFBoot2; (Hoang et al. 2018)). The local trees were summarised as a majority consensus tree with SumTrees v4.5.2, distributed with the DendroPy v4.5.2 Python library (Sukumaran and Holder 2010), with minimum clade frequency set to zero. The consensus trees were visualised using Figtree v1.4.4 (http://tree.bio.ed.ac.uk/software/figtree/).

For all trees, a specimen of *Marosatherina ladigesi* was used as an outgroup. The split between the ancestors of *M. ladigesi* and *Telmatherina* has been estimated between 12.9 and 42.9 Mya (Stelbrink et al. 2014).

#### Mitochondrial tree

We constructed a mitochondrial phylogeny from the SNP set (1836 nucleotide sites) of the mitogenomes with IQ-TREE v2.2.0 run with the GTR-model of nucleotide evolution and default parameters, using a *M. ladigesi* sample as an outgroup. The consensus tree was constructed from 1000 bootstrap trees with the UltraFast bootstrap (UFBoot2) setting in IQ-Tree (Hoang et al. 2018).

### Phylogenomic discordance

To gain insight into phylogenomic discordance in the *Telmatherina* adaptive radiation, we assessed the set of local maximum likelihood trees. We counted the number of unique topologies as well as the relative frequencies of the most frequent topologies with Newick utilities v1.6 (Junier and Zdobnov 2010). In total, there were 9150 unique topologies, meaning that every window resulted in a unique topology.

### Genomic diversity

To visualise similarity between individuals, principal component analysis (PCA) was performed on the SNP dataset with PLINK v2.0 (Purcell et al. 2007; Chang et al. 2015). For PCA analysis we only retained SNPs with a minor allele frequency (maf) of at least 0.05. PC analysis was done on the entire dataset, on the sharpfins and roundfins respectively alone, and on the sharpfins and riverine species (*T. bonti*) jointly. Figures were plotted with seaborn v 0.12.2 (Waskom 2021).

We calculated Hamming distance in the nuclear genome with PLINK v2.0 (Purcell et al. 2007; Chang et al. 2015), and divided by size of the filtered genome (accessible genome) to assess pairwise distance between all samples. We calculated heterozygosity of the entire genomes in windows of 200kb with ANGSD v0.921 (Korneliussen et al. 2014), based on the folded site frequency spectrum, using the *T. bonti* reference as the ancestral and filtering for a minimum read quality of 20, minimum mapping quality of 30 and adjusting mapping quality for excessive mismatches.

To assess whether the sharpfin, roundfin and riverine groups were genetically well separated or still shared substantial variation, we measured allele sharing across all filtered SNPs using a custom Python script with *cyvcf2* v0.31.1 (Pedersen and Quinlan 2017). For each SNP, we recorded whether both reference and alternative alleles were present within the roundfin, sharpfin, and *T. bonti* groups, as well as in every pairwise and three-way combination of these groups. We identified SNPs that were unique to each group and those shared between groups. For this analysis we did not include the presumed hybrid specimen *T. bonti 1*. Missing alleles were not considered when classifying genotypes as heterozygous. The results were visualised in a Venn diagram using *matplotlib-venn* v1.1.2. As the reference genome used for mapping and SNP calling belongs to *T. bonti*, this analysis is biased toward detecting fewer non-reference alleles in the *T. bonti* group, potentially underestimating its shared variation with the other groups. In addition, the sample sets were unbalanced across groups, which may have led to an underestimation of allele sharing between *T. bonti* (represented by fewer samples) and the other groups.

### Introgression

We calculated D-statistics (ABBA-BABA test) to assess gene flow between populations, as it allows us to distinguish between incomplete lineage sorting and introgressive hybridisation. This test compares allele sharing patterns between four groups: two sister groups (P1 and P2), a closely related third group (P3) and an outgroup (O). In the absence of gene flow, the two sister groups P1 and P2 would be expected to share the same amount of derived alleles (different from the allele in the outgroup O) with the third, related species P3. If however there is gene flow from P3 into P2, and not in P1, there will be an excess of shared derived alleles between P3 and P2, compared to P1, counted as a higher ratio of ABBA/BABA patterns. This pattern of excess allele sharing was tested for all possible (unordered) trios of individual samples in our dataset, by computing genome-wide D statistics and f4-ratio with Dsuite v0.4 (Malinsky et al. 2021). For each test, an *M. ladigesi* sample was used as an outgroup. F4-ratios of excess allele sharing between *T. bonti* and the sharpfin species relative to the roundfins is visualised, as well as excess allele sharing between *T. bonti* and the supposed hybrid specimen *T. bonti* 1.

If hybridisation events are more prevalent closer to the river-lake interface, this could lead to a spatial signal in the proportion of allele sharing of lake sharpfins with riverine *T. bonti.* We compared f4-ratios of allele sharing between *T. bonti* and sharpfins from several sampling sites at different distances from the lake-river interface, compared to a non-introgressed outgroup, the roundfins. Here we orient the roundfins as P1, the sharpfins as P2, and the riverine *T. bonti* as P3. This way, a higher f4-ratio indicates more allele sharing between *T. bonti* (P3) and the sharpfins (P2). We conducted this test for the three most common sharpfin species *T. sarasinorum*, *T. opudi* and *T. abendanoni*.

To correct for false signals caused by correlations of admixture ratios between taxa which are in the same clade as taxa involved in introgression, we calculated the f-branch statistic. This statistic takes into account the timing of introgression by investigating the internal branches of the phylogeny (Malinsky et al. 2018).

## Data availability

The genome assembly fTelBon1.1 is available on NCBI (https://www.ncbi.nlm.nih.gov/) under bioproject PRJEB51590 (https://www.ncbi.nlm.nih.gov/datasets/genome/GCA_933228915.1/)

Raw reads of the resequencing data are available at the European nucleotide archive (ENA; https://www.ebi.ac.uk/) under study PRJEB45386

## Abbreviations

GTR: General time reversible
Maf: minor allele frequency
ML: Maximum-likelihood
Mya: million years ago
NJ: Neighbour-joining

## Acknowledgements

No samples have been collected for this study. The specimens used in this study were available from museum collections of the Museum Zoologicum Bogoriense (MZB) in Bogor, Indonesia and the Leibniz Institute for the Analysis of Biodiversity Change (LIB), Bonn, Germany. Original sampling was conducted between 2002 and 2010 with permits from the Indonesian government (Surat Izin Penelitian) 5602/SU/KS/2002 (LIPI) and 0093/SIP/FRP/SM/V/2010 (RISTEK). We thank the Heads of Research Center for Biology, Indonesian Institute of Sciences (LIPI; presently Research Center for Biosystematics and Evolution, National Research and Innovation Agency or Badan Riset dan Inovasi Nasional [BRIN] and the Kementerian Negara Riset dan Teknologi [RISTEK] presently also BRIN) for permissions to conduct research in Indonesia. The late R. K. Hadiaty enabled joint fieldwork in Sulawesi, funded by the Deutsche Forschungsgemeinschaft (DFG SCHL 567/2, HE 5707/2-1, HE 5707/3-1), providing voucher samples re-analyzed in the present project. Computational resources and services used in this work were provided by the VSC (Flemish Supercomputer Center), funded by the Research Foundation - Flanders (FWO) and the Flemish Government. Access to VSC infrastructure was facilitated by the CalcUA team at Universiteit Antwerpen. ELRDK is supported as postdoctoral fellow fundamental research by the Flemish Research Foundation (FWO postdoctoral scholarship 1275922N). Research and collaboration between Indonesia and Europe was supported by Research Foundation - Flanders (FWO) (project G0A9B24N to HS and travel grant V408023N to ELRDK), VLIR-UOS (SI project ID2022SIN349A102 to HS and GMSR project 54084 to ELRDK) and Universiteit Antwerpen (BOF Seal of approval and KPBOF project 48620 to ELRDK).

## Author contributions

ELRDK, FH, AB and HS conceived the study. ELRDK, CCJ, VB and HS conducted analysis on resequencing data. SK, AD and GO conducted molecular laboratory work. The Wellcome Sanger Institute Tree of life programme organised and conducted fTelBon1.1 reference genome assembly and annotation. AT led manual curation of the fTelBon1.1 reference genome. DW, DFM and FH organised and conducted data collection. ELRDK and HS wrote the manuscript with input from all co-authors.

