## Supplementary materials for "Ancient and recent riverine gene flow contributed to the adaptive radiation of sailfin silversides in Wallace’s Dreampond"

### Multiple instances of river–lake introgression in the adaptive radiation of sailfin silversides in Wallace's Dreampond

[4] members: https://doi.org/10.5281/zenodo.4783585

[5] members: https://doi.org/10.5281/zenodo.4790455

[6] members: https://doi.org/10.5281/zenodo.7116866

[7] Museum Zoologicum Bogoriense, Research Center for Biosystematics and Evolution, National Research and Innovation Agency (BRIN), Cibinong, 16911, Indonesia

[8] Naturalis Biodiversity Center, 2333 Leiden, The Netherlands

Corresponding author: Els L. R. De Keyzer

#### Supplementary Materials

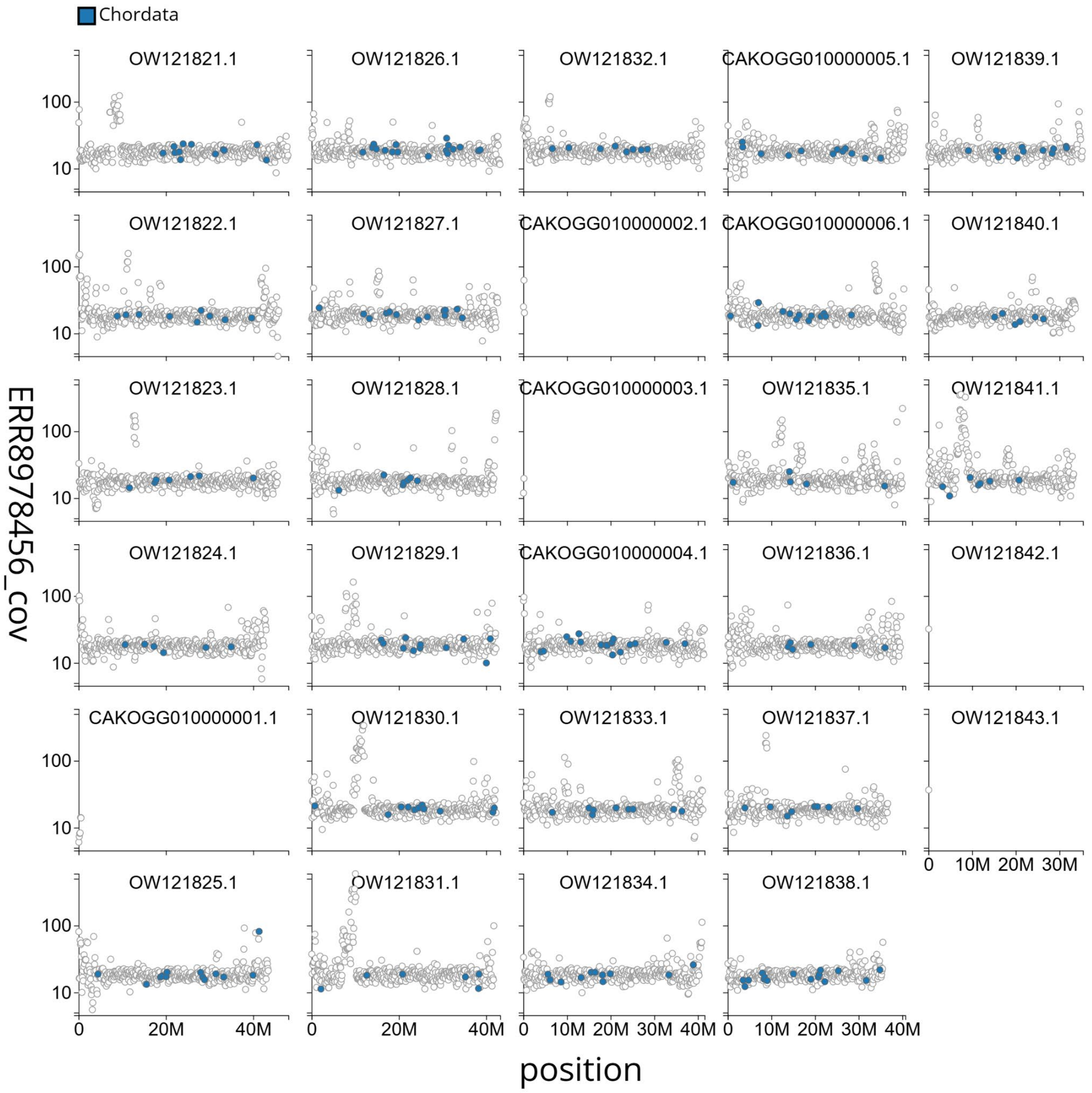


Supplementary figure S1. **Distribution plot of base coverage in ERR8978456 against position for sequences in assembly CAKOGG01.** 100kb windows are coloured by phylum.

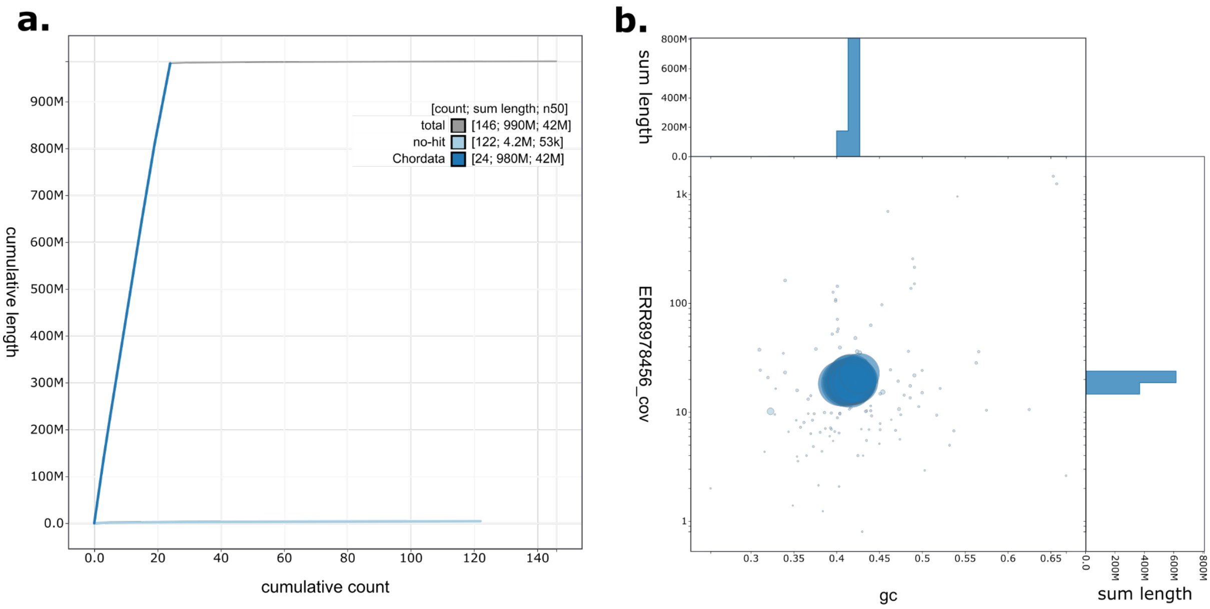


supplementary figure S2. **a. Cumulative sequence length for assembly CAKOGG01.** The grey line shows cumulative length for all sequences. Coloured lines show cumulative lengths of sequences assigned to each phylum using the buscogenes taxrules. An interactive version of this figure is available at https://blobtoolkit.genomehubs.org/view/Telmatherina%20bonti/dataset/CAKOGG01/cumulative **b. Blob plot of base coverage in ERR8978456 against GC proportion for sequences in assembly CAKOGG01.** Sequences are coloured by phylum. Circles are sized in proportion to sequence length. Histograms show the distribution of sequence length sum along each axis. An inveractive version of this figure is available at https://blobtoolkit.genomehubs.org/view/Telmatherina%20bonti/dataset/CAKOGG01/blob

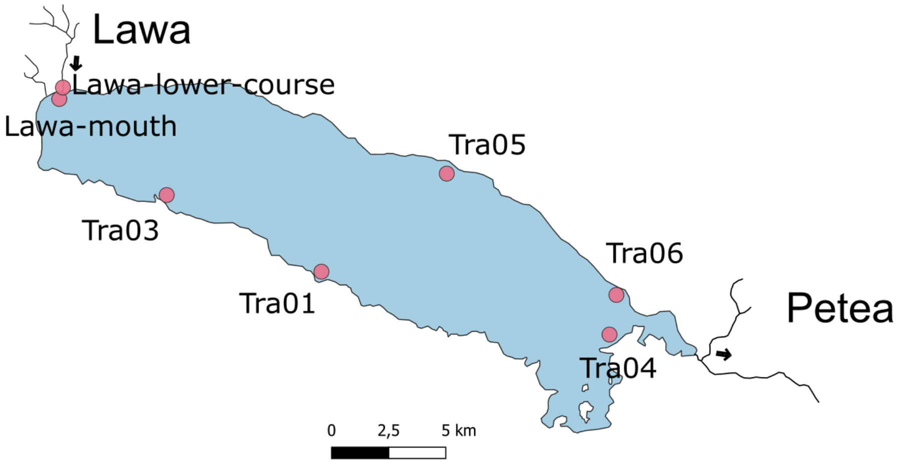


Supplementary figure S3. **Map of sampling locations.** Names of sampling locations correspond to sample information as displayed in supplementary table S2. Inflow river Lawa and outflow river Petea are indicated.

**a.**

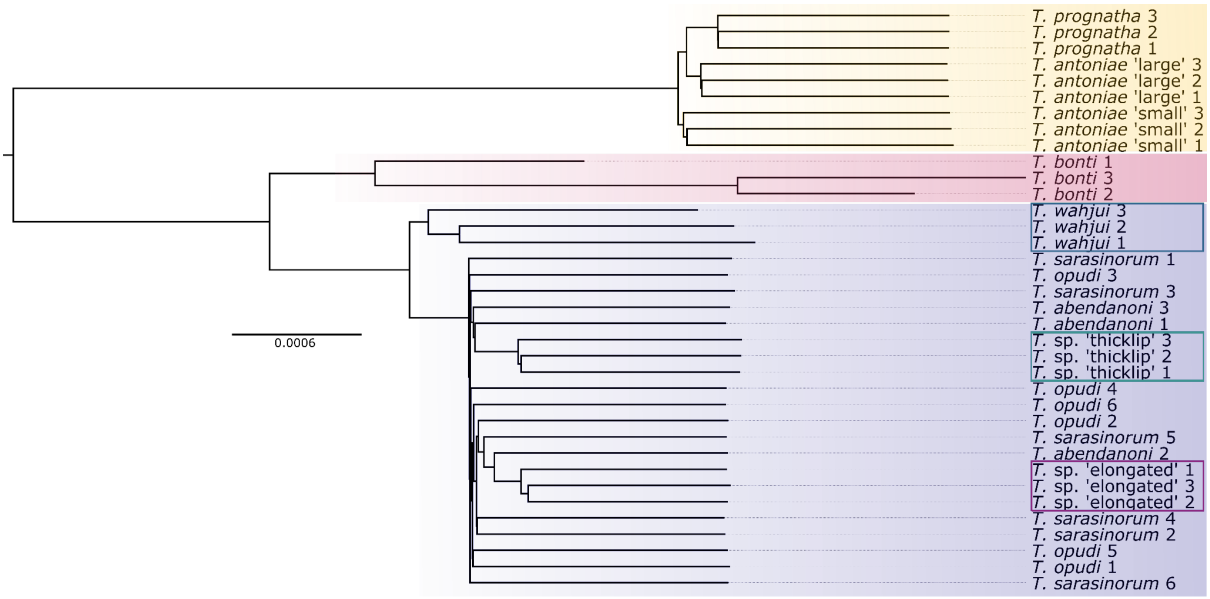


**b.**

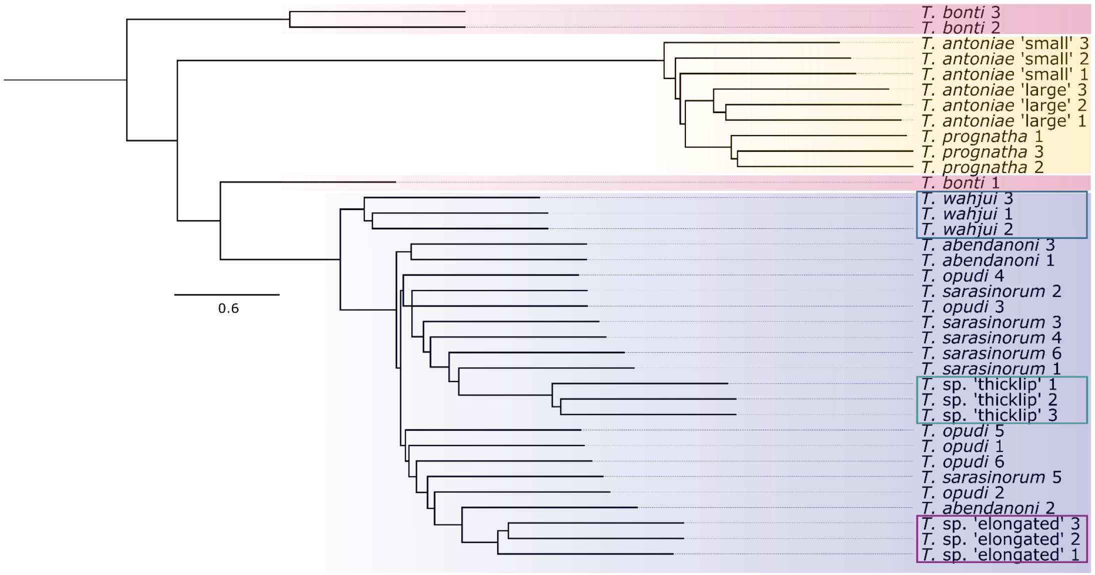


Supplementary figure S4. a. **NJ tree based on distance matrix**. Scale is the number of mutations divided by accessible genome size b. **astral consensus of NJ trees** based local distance trees based on 100 kb windows. All trees are rooted against a sample of the outgroup species *M. ladigesi*.

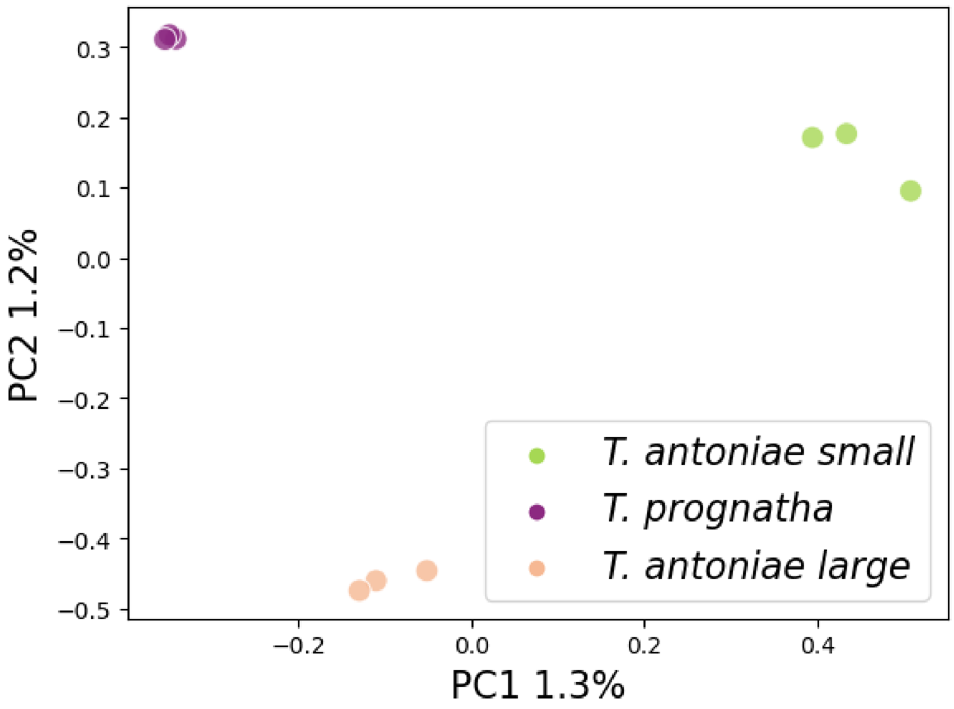


Supplementary figure S5. **Visualisation of the first and second principal components** based on the SNP dataset for Lake Matano roundfin species.

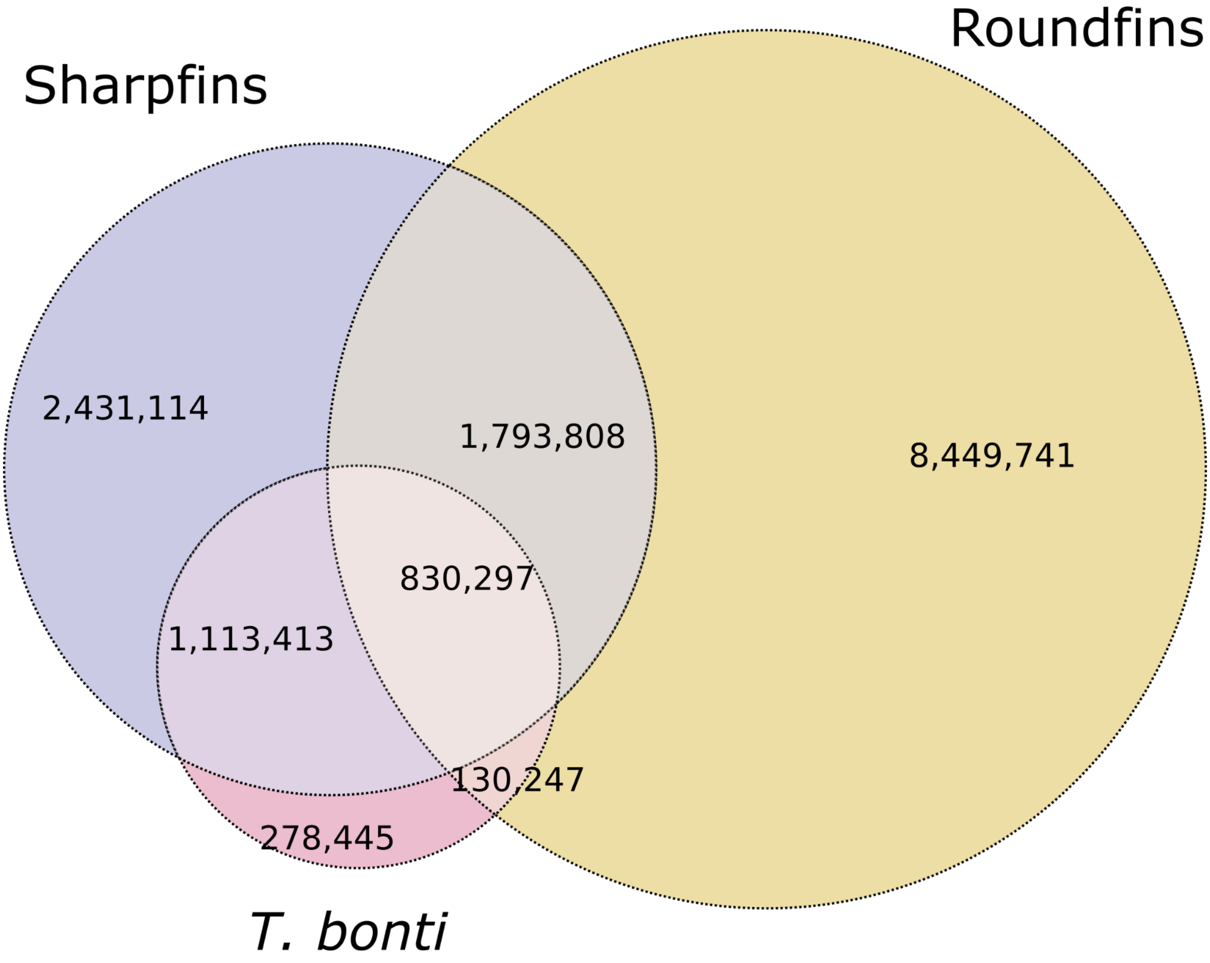


Supplementary figure S6. **Venn-diagram of shared and private variable alleles** in sharpfins (n = 24), roundfins (n = 9) and *T. bonti* (n = 2). Outer circles represent the number of variable alleles unique to the corresponding group. Overlapping areas indicate the number of variable alleles shared between the respective groups.

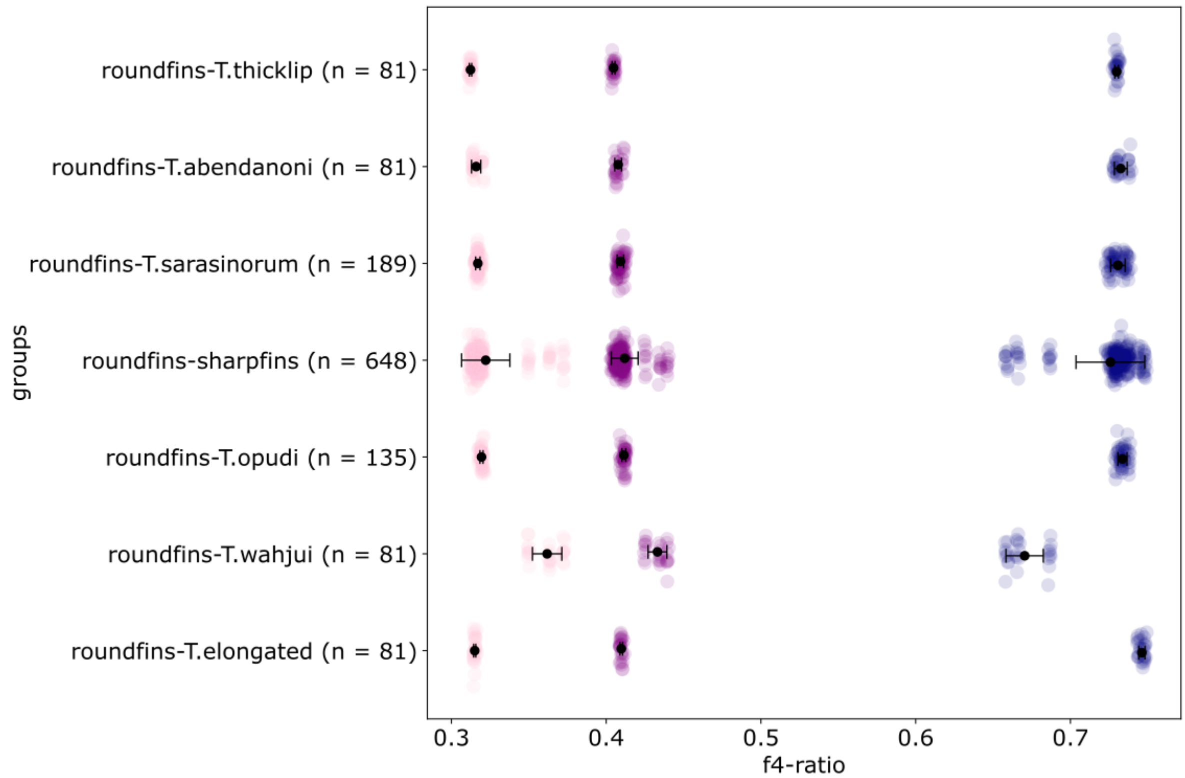


Supplementary figure S7. **Excess allele sharing (f4-ratio)** between *T. bonti* 3 (pink), *T. bonti* 2 (purple), *T. bonti* 1 (blue) (P3) and the sharpfin species indicated on Y axis (P2), relative to the roundfins (P1)

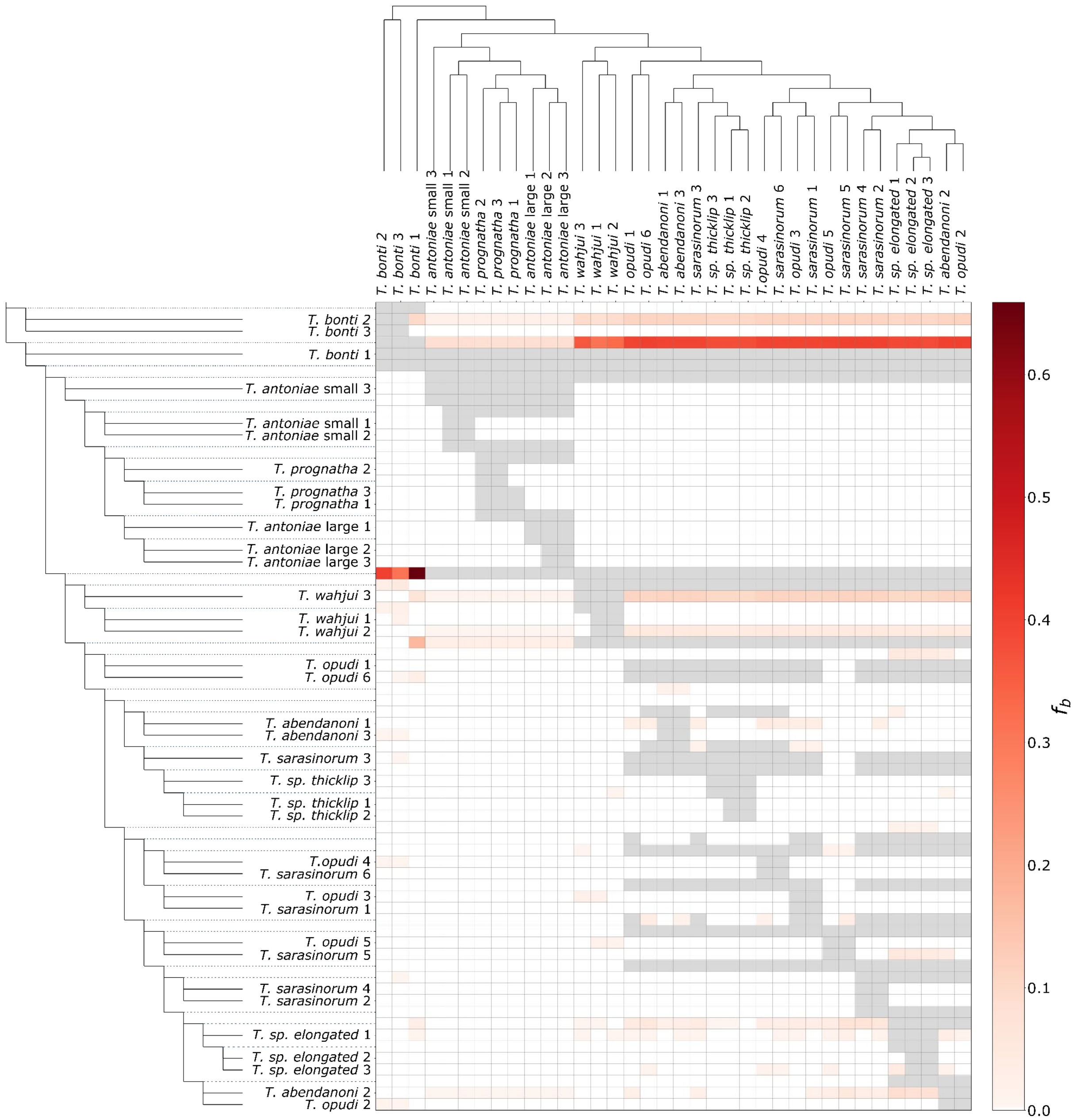


Supplementary figure S8. **Heatmap of pairwise f-branch (fb) statistic**, based on SNP dataset. Higher values (darker colour) indicate more excess allele sharing between specimens.

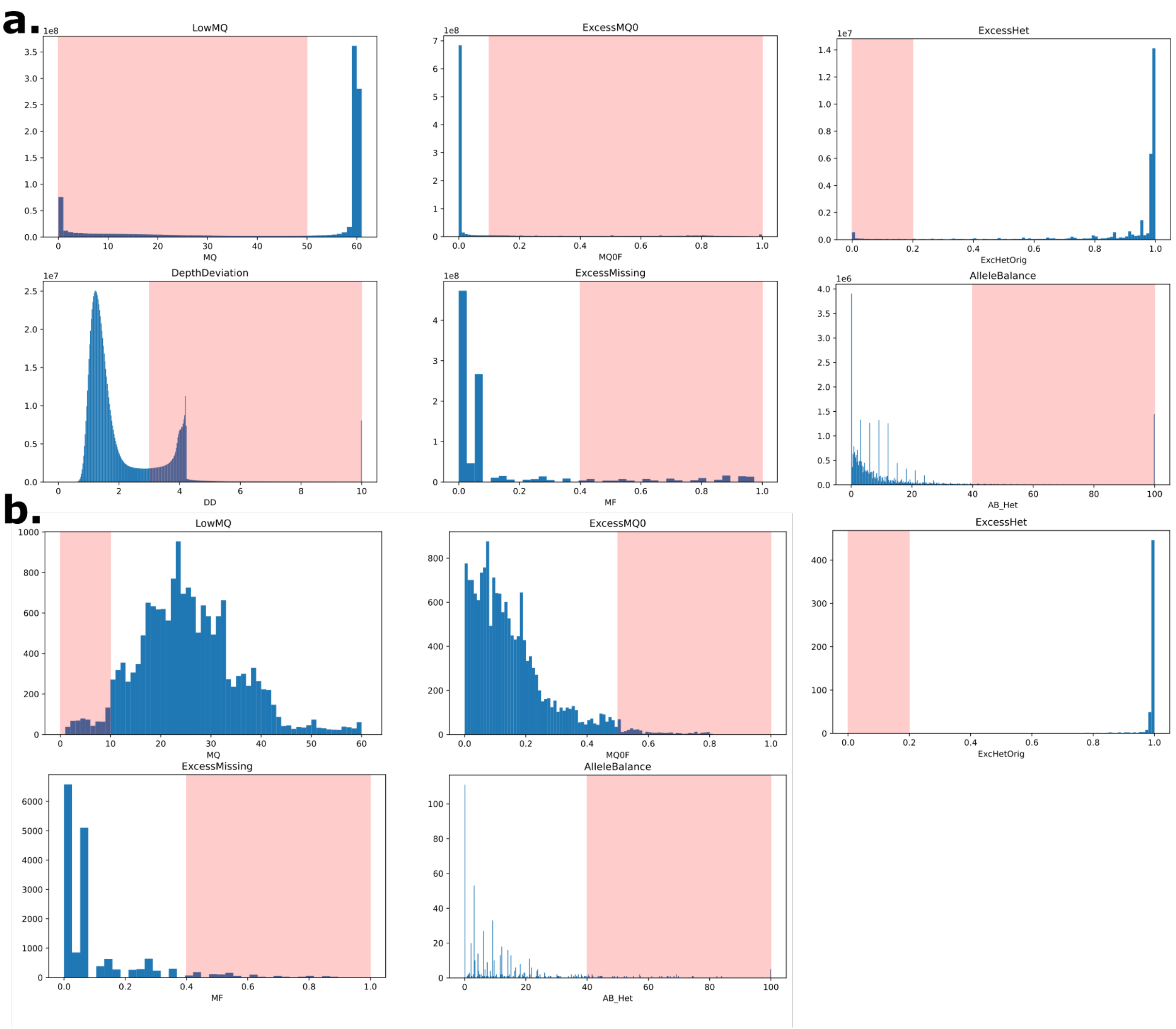


Supplementary figure S9. **Filter settings and number of sequences removed per filter**. Area highlighted in red indicates the filter threshold (a) for the nuclear genome and (b) for the mitochondrial DNA.

Supplementary table S1. **Chromosomes of fTelBon1:** chromosome size, GC-content in percentage, average coverage and accession number

| Chromosome | Size (Mb) | GC% | Coverage | Accession |
|---|---|---|---|---|
| 1 | 48.06 | 0.41 | 18.53 | OW121821.1 |
| 2 | 45.92 | 0.417 | 19.52 | OW121822.1 |
| 3 | 45.68 | 0.416 | 18.00 | OW121823.1 |
| 4 | 43.59 | 0.419 | 18.72 | OW121824.1 |
| 5 | 43.20 | 0.418 | 18.68 | OW121825.1 |
| 6 | 43.05 | 0.405 | 18.30 | OW121826.1 |
| 7 | 42.78 | 0.414 | 19.30 | OW121827.1 |
| 8 | 42.41 | 0.415 | 18.59 | OW121828.1 |
| 9 | 42.36 | 0.416 | 19.31 | OW121829.1 |
| 10 | 42.35 | 0.417 | 21.26 | OW121830.1 |
| 11 | 41.80 | 0.415 | 20.91 | OW121831.1 |
| 12 | 41.52 | 0.41 | 18.73 | OW121832.1 |
| 13 | 41.29 | 0.412 | 18.33 | OW121833.1 |
| 14 | 41.02 | 0.421 | 19.99 | OW121834.1 |
| 15 | 40.93 | 0.415 | 18.58 | OW121835.1 |
| 16 | 40.68 | 0.423 | 19.09 | OW121836.1 |
| 17 | 40.41 | 0.419 | 19.10 | OW121837.1 |
| 18 | 40.13 | 0.419 | 20.43 | OW121838.1 |
| 19 | 39.80 | 0.421 | 19.07 | OW121839.1 |
| 20 | 36.75 | 0.422 | 18.81 | OW121840.1 |
| 21 | 35.62 | 0.416 | 18.23 | OW121841.1 |
| 22 | 35.39 | 0.414 | 18.52 | OW121842.1 |
| 23 | 33.76 | 0.419 | 18.54 | OW121843.1 |
| 24 | 33.33 | 0.427 | 22.75 | OW121844.1 |

Supplementary table S2. **sample information -** sample ID as used in the manuscript and additional information about the samples used in the study. Sampling locations correspond to locations indicated in Supplementary figure S3.

| individual_id | species | group | sex | sampling_date | catchment | sampling_location | ZFMK_collection_ID | analysis_id | Latitude | Longitude |
|---|---|---|---|---|---|---|---|---|---|---|
| *T. bonti 2* | *Telmatherina bonti* | T. bonti | M | 18/11/2006 | Lake Matano | Lawa-lower-course | 4360 | BO4360 | -2.429848 | 121.224196 |
| *T. bonti 3* | *Telmatherina bonti* | T. bonti | M | 18/11/2006 | Lake Matano | Lawa-lower-course | 4384 | BO4384 | -2.429848 | 121.224196 |
| *T. opudi 6* | *Telmatherina opudi* | sharpfin | M | 18/11/2006 | Lake Matano | Lawa-lower-course | 4365 | BO4365 | -2.429848 | 121.224196 |
| *T. sarasinorum 4* | *Telmatherina sarasinorum* | sharpfin | M | 18/11/2006 | Lake Matano | Lawa-lower-course | 4364 | SA4364 | -2.429848 | 121.224196 |
| *T. bonti 1* | *Telmatherina bonti* | T. bonti | F | 14/11/2006 | Lake Matano | Lawa-mouth | 4164 | BO4164 | -2.431242 | 121.222230 |
| *T. opudi 1* | *Telmatherina opudi* | sharpfin | M | 19/11/2006 | Lake Matano | Lawa-mouth | 4396 | OP4396 | -2.431242 | 121.222230 |
| *T. opudi 2* | *Telmatherina opudi* | sharpfin | M | 19/11/2006 | Lake Matano | Lawa-mouth | 4397 | OP4397 | -2.431242 | 121.222230 |
| *T. sarasinorum 2* | *Telmatherina sarasinorum* | sharpfin | M | 14/11/2006 | Lake Matano | Lawa-mouth | 4094 | SA4094 | -2.431242 | 121.222230 |
| *T. sarasinorum 3* | *Telmatherina sarasinorum* | sharpfin | M | 11/12/2006 | Lake Matano | Lawa-mouth | 4324 | SA4324 | -2.431242 | 121.222230 |
| *T. sarasinorum 5* | *Telmatherina sarasinorum* | sharpfin | M | 14/11/2006 | Lake Matano | Lawa-mouth | 4095 | SA4095 | -2.431242 | 121.222230 |
| *T. abendanoni 3* | *Telmatherina abendanoni* | sharpfin | M | 18/11/2002 | Lake Matano | Tra01 | 1831 | AB1831 | -2.506090 | 121.326814 |
| *T. antoniae* 'large' *1* | *Telmatherina antoniae* 'large' | roundfin | M | 20/10/2002 | Lake Matano | Tra01 | 225 | AL225 | -2.506090 | 121.326814 |
| *T. antoniae* 'large' *2* | *Telmatherina antoniae* 'large' | roundfin | M | 20/10/2002 | Lake Matano | Tra01 | 226 | AL226 | -2.506090 | 121.326814 |
| *T. antoniae* 'large' *3* | *Telmatherina antoniae* 'large' | roundfin | F | 20/10/2002 | Lake Matano | Tra01 | 233 | AL233 | -2.506090 | 121.326814 |
| *T. antoniae* 'small' *1* | *Telmatherina antoniae* 'small' | roundfin | M | 15/10/2002 | Lake Matano | Tra01 | 129 | AS129 | -2.506090 | 121.326814 |
| *T. antoniae* 'small' *2* | *Telmatherina antoniae* 'small' | roundfin | F | 15/10/2002 | Lake Matano | Tra01 | 131 | AS131 | -2.506090 | 121.326814 |
| *T. prognatha 1* | *Telmatherina prognatha* | roundfin | M | 20/10/2002 | Lake Matano | Tra01 | 216 | PR216 | -2.506090 | 121.326814 |
| *T. prognatha 2* | *Telmatherina prognatha* | roundfin | F | 20/10/2002 | Lake Matano | Tra01 | 222 | PR222 | -2.506090 | 121.326814 |
| *T. prognatha 3* | *Telmatherina prognatha* | roundfin | M | 04/11/2002 | Lake Matano | Tra01 | 444 | PR444 | -2.506090 | 121.326814 |
| *T. antoniae* 'small' *3* | *Telmatherina antoniae* 'small' | roundfin | M | 31/10/2002 | Lake Matano | Tra03 | 330 | AS330 | -2.474489 | 121.263214 |
| *T. abendanoni 1* | *Telmatherina abendanoni* | sharpfin | M | 14/11/2002 | Lake Matano | Tra04 | 1224 | AB1224 | -2.529604 | 121.444242 |
| *T. wahjui 1* | *Telmatherina wahjui* | sharpfin | M | 14/11/2002 | Lake Matano | Tra04 | 1238 | WA1238 | -2.529604 | 121.444242 |
| *T. wahjui 2* | *Telmatherina wahjui* | sharpfin | F | 14/11/2002 | Lake Matano | Tra04 | 1436 | WA1436 | -2.529604 | 121.444242 |
| *T. wahjui 3* | *Telmatherina wahjui* | sharpfin | M | 14/11/2002 | Lake Matano | Tra04 | 1315 | WA1315 | -2.529604 | 121.444242 |
| *T. sp. elongated 3* | *Telmatherina* sp. 'elongated' | sharpfin | F | 14/11/2002 | Lake Matano | Tra04 | 1325 | EY1325 | -2.529604 | 121.444242 |
| *T.* sp. 'thicklip' *1* | *Telmatherina* sp. 'thicklip' | sharpfin | M | 14/11/2002 | Lake Matano | Tra04 | 1240 | TL1240 | -2.529604 | 121.444242 |
| *T.* sp. 'thicklip' *2* | *Telmatherina* sp. 'thicklip' | sharpfin | M | 14/11/2002 | Lake Matano | Tra04 | 1309 | TL1309 | -2.529604 | 121.444242 |
| *T. opudi 4* | *Telmatherina opudi* | sharpfin | M | 08/11/2002 | Lake Matano | Tra05 | 719 | OP719 | -2.460722 | 121.380139 |
| *T. opudi 5* | *Telmatherina opudi* | sharpfin | F | 08/11/2002 | Lake Matano | Tra05 | 743 | OP743 | -2.460722 | 121.380139 |
| *T. sarasinorum 1* | *Telmatherina sarasinorum* | sharpfin | M | 29/06/2010 | Lake Matano | Tra05 | 8719 | SA8719 | -2.460722 | 121.380139 |
| *T. opudi 3* | *Telmatherina sarasinorum* | sharpfin | M | 29/06/2010 | Lake Matano | Tra05 | 8717 | OP8717 | -2.460722 | 121.380139 |
| *T. sarasinorum 6* | *Telmatherina sarasinorum* | sharpfin | M | 29/06/2010 | Lake Matano | Tra05 | 8715 | SA8715 | -2.460722 | 121.380139 |
| *T.* sp. 'thicklip' *3* | *Telmatherina* sp. 'thicklip' | sharpfin | F | 29/06/2010 | Lake Matano | Tra05 | 8714 | TL8714 | -2.460722 | 121.380139 |
| *T. abendanoni 2* | *Telmatherina abendanoni* | sharpfin | M | 26/11/2002 | Lake Matano | Tra06 | 2013 | AB2013 | -2.509969 | 121.445643 |
| *T.* sp. 'elongated' *1* | *Telmatherina* sp. 'elongated' | sharpfin | F | 26/11/2002 | Lake Matano | Tra06 | 2015 | EL2015 | -2.509969 | 121.445643 |
| *T.* sp. 'elongated' *2* | *Telmatherina* sp. 'elongated' | sharpfin | M | 26/11/2002 | Lake Matano | Tra06 | 2016 | EL2016 | -2.509969 | 121.445643 |
| *M. ladigesi 1* | *Marosatherina ladigesi* | outgroup | M | | | ZFMK aquarium | | MP3 | | |
